# Resident memory CD8 T cells persist for years in human small intestine

**DOI:** 10.1101/553792

**Authors:** Raquel Bartolomé Casado, Ole J.B. Landsverk, Sudhir Kumar Chauhan, Lisa Richter, Dang Phung, Victor Greiff, Louise Fremgaard Risnes, Ying Yao, Ralf Stefan Neumann, Sheraz Yaqub, Ole Øyen, Rune Horneland, Einar Martin Aandahl, Vemund Paulsen, Ludvig M. Sollid, Shuo-Wang Qiao, Espen S. Bækkevold, Frode L. Jahnsen

## Abstract

In human small intestine, most CD8 T cells in the lamina propria and epithelium express a resident memory (Trm) phenotype and persist for at least one year in transplanted tissue. Intestinal CD8 Trm cells have a clonally expanded immune repertoire that is stable over time and exhibit enhanced protective capabilities.

**Graphical abstract:** 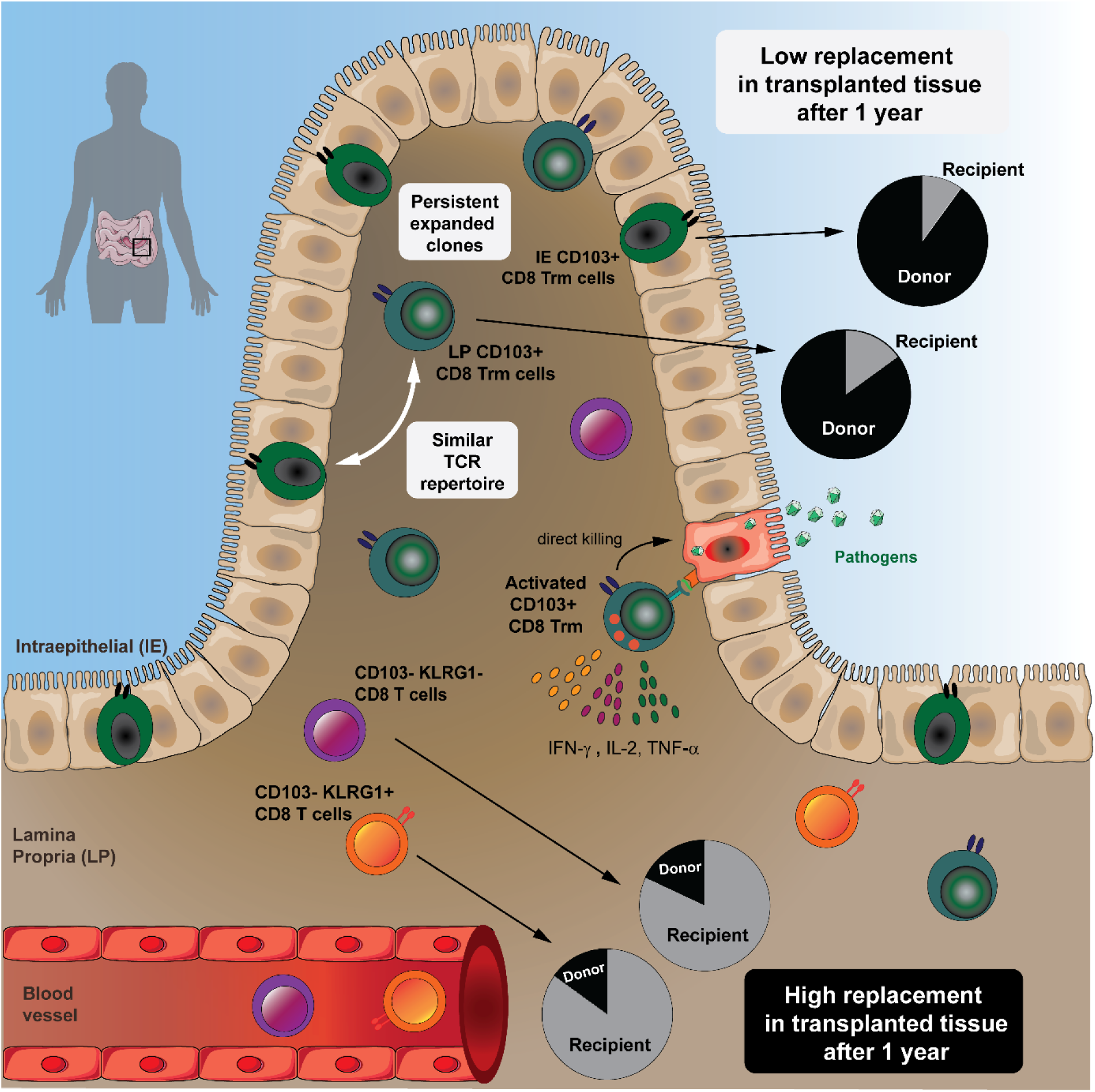

**Highlights:**

- The vast majority of CD8 T cells in the human small intestine are Trm cells
- CD8 Trm cells persist for >1 year in transplanted duodenum
- Intraepithelial and lamina propria CD8 Trm cells show highly similar TCR repertoire
- Intestinal CD8 Trm cells efficiently produce cytokines and cytotoxic mediators

**Abbreviations:**

IE: intraepithelial
LP: lamina propria
RPMI: Roswell Park Memorial Institute medium
SI: small intestine
TCR: T cell receptor
Trm: resident memory T cell
Tx: pancreatic-duodenal transplantation (of diabetes mellitus patients)

## Summary

Resident memory CD8 T cells (Trm) have been shown to provide effective protective responses in the small intestine (SI) in mice. A better understanding of the generation and persistence of SI CD8 Trm cells in humans may have implications for intestinal immune-mediated diseases and vaccine development. Analyzing normal and transplanted human SI we demonstrated that the majority of SI CD8 T cells were *bona fide* CD8 Trm cells that survived for over 1 year in the graft. Intraepithelial and lamina propria CD8 Trm cells showed a high clonal overlap and a repertoire dominated by expanded clones, conserved both spatially in the intestine and over time. Functionally, lamina propria CD8 Trm cells were potent cytokine-producers, exhibiting a polyfunctional (IFN-γ+ IL-2+ TNF-α+) profile, and efficiently expressed cytotoxic mediators after stimulation. These results suggest that SI CD8 Trm cells could be relevant targets for future oral vaccines and therapeutic strategies for gut disorders.

## Introduction

Studies in mice have shown that the intestine contains high numbers of resident memory CD8 T (Trm) cells (Steinert et al., 2015). CD8 Trm cells are persistent, non-circulatory cells, which provide particularly rapid and efficient protection against recurrent infections (Ariotti et al., 2014). In addition to their cytotoxic activity, CD8 Trm cells are efficient producers of pro-inflammatory cytokines that rapidly trigger both innate and adaptive protective immune responses (Schenkel et al., 2014). Thus, CD8 Trm cells are attractive targets for vaccine development against intracellular pathogens (Gola et al., 2018) and cancer immunotherapy (Park et al., 2018).

Although studies in mice have significantly advanced our understanding of CD8 Trm cell function in the intestine (Jabri and Ebert, 2007; Konjar et al., 2018; Sheridan et al., 2014), translation of mouse data into humans should be implemented with caution. Specifically, the generation of T cell memory in humans occurs after exposure to a broad variety of pathogens and commensals over many decades of life, which cannot be recapitulated in mouse models. Most mice are maintained in specific pathogen-free (SPF) conditions that reduce their microbiome diversity, which in turn influence the immune homeostasis and response to pathogens in the intestine (Maynard et al., 2012; Tao and Reese, 2017). Moreover, murine intraepithelial (IE) CD8 T cells constitute a heterogeneous population of unconventional CD8αα+ and conventional CD8αβ+ T cells, with partially overlapping effector properties but different developmental origins (McDonald et al., 2018). In contrast, their human counterparts mainly consist of conventional CD8αβ-expressing cells (Jabri and Ebert, 2007).

Under steady state conditions the human small intestine (SI) is densely populated by CD8 T cells, both in the lamina propria (LP) and within the epithelium. However, most studies of human SI CD8 T cells have focused exclusively on intraepithelial (IE) CD8 T cells (Abadie et al., 2012). Therefore, there is currently very limited knowledge about several functional aspects of SI CD8 T cells. For example, is the CD8 T-cell population heterogeneous? Are the CD8 T cells persistent or circulating cells? What are their cytotoxic and cytokine-producing capacity? To what extent are LP and IE CD8 T cells clonally related? In order to consider using SI CD8 T cells as targets to design effective oral vaccines and to understand their involvement in immune-mediated diseases these are critical questions to address. Analyzing the CD8 T cell compartment in the normal SI as well as in a unique transplantation setting in humans, we report that most SI CD8 T cells are Trm cells that persist for more than a year in the epithelium and LP. High-throughput T cell receptor (TCR) sequencing showed a polarized immune repertoire and high clonal relatedness between IE and LP CD8 Trm cells. Functionally, the minor population of LP CD103-CD8 T cells was transient in the tissue and produced less cytokines but had more pre-formed cytotoxic granules. In contrast, we demonstrate that activated CD8 Trm cells expressed high levels of multiple cytokines (polyfunctional) and showed *de novo* production of cytotoxic granules. Thus, SI CD8 Trm have potent protective capabilities and present functional roles similar to CD8 Trm described in mice.

## Results

### Most CD8 T cells in the human small intestine express a resident memory phenotype

To determine whether the human SI contains persisting CD8 T cells, we first examined biopsies from donor duodenum after pancreatic-duodenal transplantation (Tx) of type I diabetic patients (Horneland et al., 2015). Using fluorescent *in situ* hybridization probes specific for X/Y-chromosomes on tissue sections where patient and donor were of different gender, we could precisely distinguish donor and recipient cells, and consistently detected a high fraction of persisting donor CD8 T cells one year after transplantation **(Figure 1A).**

**Figure 1.**
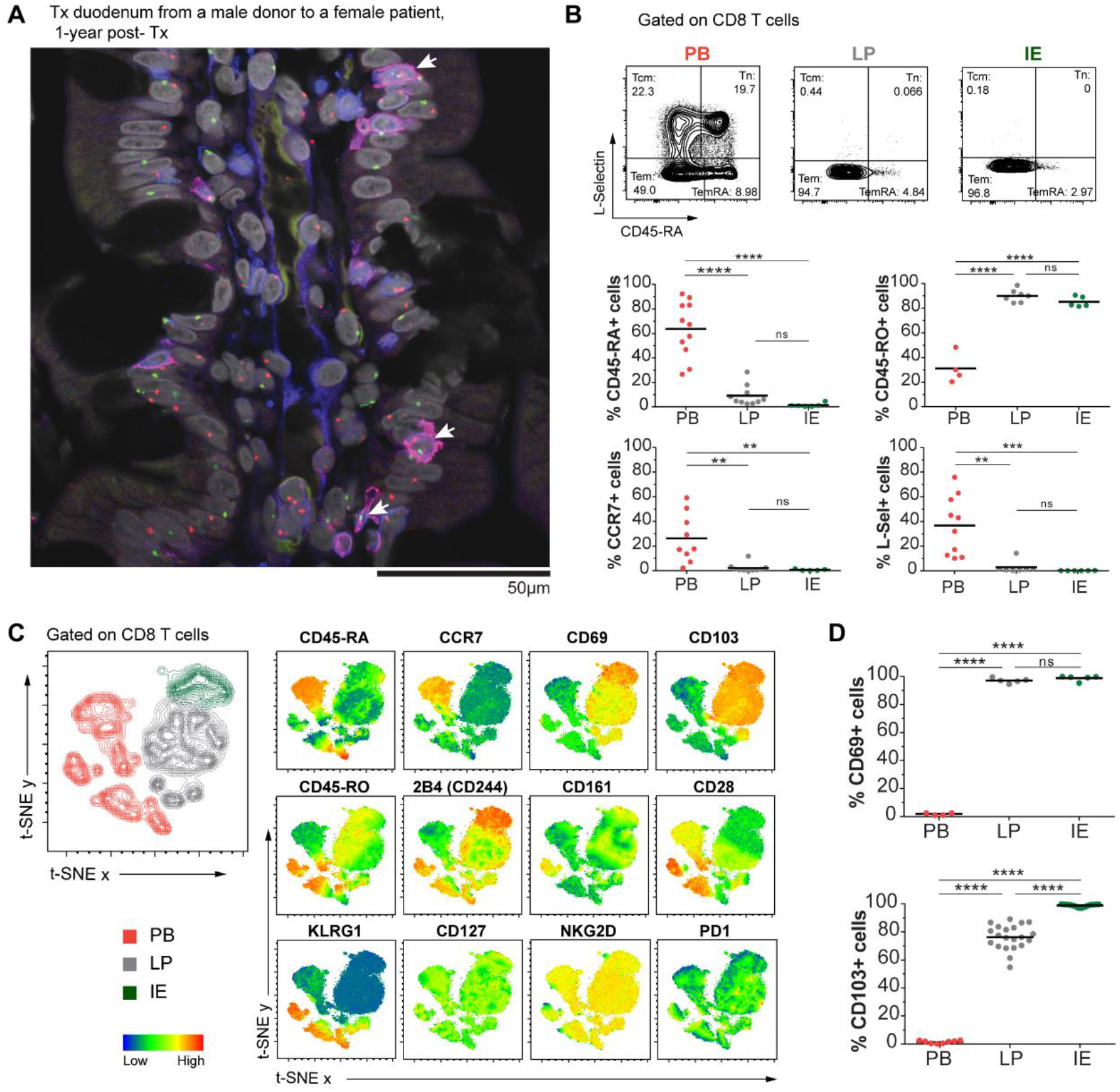
Human small intestinal mucosa harbors a substantial population of CD8 T cells with a Trm phenotype. (**A**). Representative confocal image of a tissue section from a male donor duodenum one year after transplantation into a female patient stained with X/Y chromosome fluorescent *in situ* hybridization probes (Y, green; X, red) and antibodies against CD8 (red) and CD3 (blue). Hoechst (gray) stains individual nuclei and white arrows indicate donor (male) CD8 T cells (n=8). (**B**) Expression of classical lymph node homing and memory markers on peripheral blood (PB), intestinal lamina propria (LP) and intraepithelial (IE) CD8 T cells. Representative contour-plots and compiled data for each marker are given. (**C**) t-SNE map showing the distribution of PB (red), LP (grey) and IE (green) CD8 T cell clusters (left). Overlay of the t-SNE map with expression levels for each marker, color-coded based on the median fluorescent intensity values representative of 3 samples. See **Figure S3** for details on preprocessing of flow data for t-SNE analysis. (**D**) Compiled data for the expression of CD69 and CD103 on PB, LP and IE CD8 T cells. Black bars in (B, D) indicate mean value. Statistical analysis was performed using one-way ANOVA with Tukey’s multiple comparisons test. ns, not significant; *, p≤0.05; **, p≤0.01; ***, p≤0.001; ****, p≤0.0001.

This finding encouraged us to further characterize SI CD8 T cells, and to this end we examined resections of proximal SI obtained from pancreatic cancer surgery (Whipple procedure), and from donors and recipients during pancreatic-duodenal Tx (baseline samples). All tissue samples were evaluated by pathologists and only histologically normal SI was included. Peripheral blood (PB) was collected from Tx recipients and pancreatic cancer patients, and mononuclear cells (PBMCs) were isolated and analyzed by flow cytometry together with single-cell suspensions from enzyme-digested LP and IE. The cross-contamination between IE and LP fractions was negligible, attested by complete removal of epithelial cells assessed by histology (**Figure S1A**) and by low numbers of epithelial cells in the LP fraction detected by flow cytometry (**Figure S1B-C**). We found that, in blood and SI-LP, CD8 T cells constituted about a third of the CD3 T cells, while CD8 T cells dominated (>75%) in the epithelium **(Figure S2A)**. Almost all SI CD8 T cells were TCRαβ+ (∼99.8% in LP and 98.7% in epithelium, **Figure S2B**), and virtually all expressed the co-receptor CD8αβ **(Figure S2C**). The vast majority of SI CD8 T cells exhibited a CD45RO+ CD45RA-L-Sel-CCR7-effector memory (Tem) phenotype **(Figure 1B),** whereas PB also contained a substantial fraction of naïve (Tn, CD45RO-CD45RA+ CCR7+ L-Sel+) and central memory (Tcm, CD45RO+ CD45RA-CCR7+ L-Sel+) CD8 T cells (**Figure 1B**).

To further define and compare LP and IE CD8 T cells with PB CD8 T cells, we performed t-SNE analysis including classical Trm markers and functional (e.g. costimulatory and inhibitory) receptors associated with a Trm phenotype (Fergusson et al., 2016; Kumar et al., 2017; Mackay et al., 2013; Thome et al., 2014). LP and IE CD8 T cells clustered together and were distinct from PB CD8 T cells (**Figure 1C, left**). By visualizing the expression pattern for each marker, we confirmed that cells expressing CD45RA and CCR7 were confined to the PB clusters, whereas the Trm markers CD69 and CD103 were only expressed on tissue-derived CD8 T cells. SI CD8 T cells also showed high expression of the co-inhibitory receptor 2B4 (SLAMF4) and the C-type lectin NK receptor CD161, whereas CD28 and killer-cell lectin like receptor G1 (KLRG1) were expressed at higher levels on PB CD8 T cells. CD127 (IL7 receptor-α), NKG2D and PD-1 did not show any clustered expression (**Figure 1C**).

Virtually all LP and IE CD8 T cells expressed CD69 (97% and 99.5%, respectively). CD103 was expressed by most (76%) LP CD8 T cells and by all IE CD8 T cells (**Figure 1D**), whereas the minor population of LP CD8 T cells lacking CD103 was more similar to PB CD8 T cells (**Figure 1C**). Given that CD103 is the currently most established marker to infer residency at mucosal surfaces (Beura et al., 2018b) we evaluated the phenotypic profiles of CD103+ and CD103-SI CD8 T cells in a larger cohort of donors **(Figure 2A)**. CD127 was expressed by a larger fraction of LP CD103+ CD8 T cells than IE CD103+ and LP CD103-CD8 T cells **(Figure 2A).** However, IE CD103+ CD8 T cells had higher expression of 2B4 and CD161 compared to CD103-CD8 T cells. NKG2D was broadly expressed on all SI CD8 T cell subsets, but showed significantly higher expression in the CD103-compartment. In addition, we found consistently higher numbers of cells positive for CD28, PD-1 and KLRG1 within the CD103-CD8 T population compared to both LP and IE CD103+ cells **(Figure 2A)**. Interestingly, around half of the CD103-CD8 T cells expressed KLRG1, whereas most CD103+ CD8 T cells were KLRG1-, and plotting CD103 *versus* KLRG1 divided LP CD8 T cells into three distinct subsets **(Figure 2B).** The distribution of these three LP CD8 T cell subsets was conserved lengthwise in the SI, as demonstrated by their similar representation in mucosal biopsies taken several cm apart the same SI **(Figure S4)**. We also analyzed the expression of the proliferation marker Ki67 by intracellular staining and flow cytometry. We detected few Ki67 positive cells (median <5%) in all CD8 T cell subsets from normal SI samples **(Figure 2C)**, indicating that the proliferative activity of SI CD8 T cells under steady-state conditions is low.

**Figure 2.**
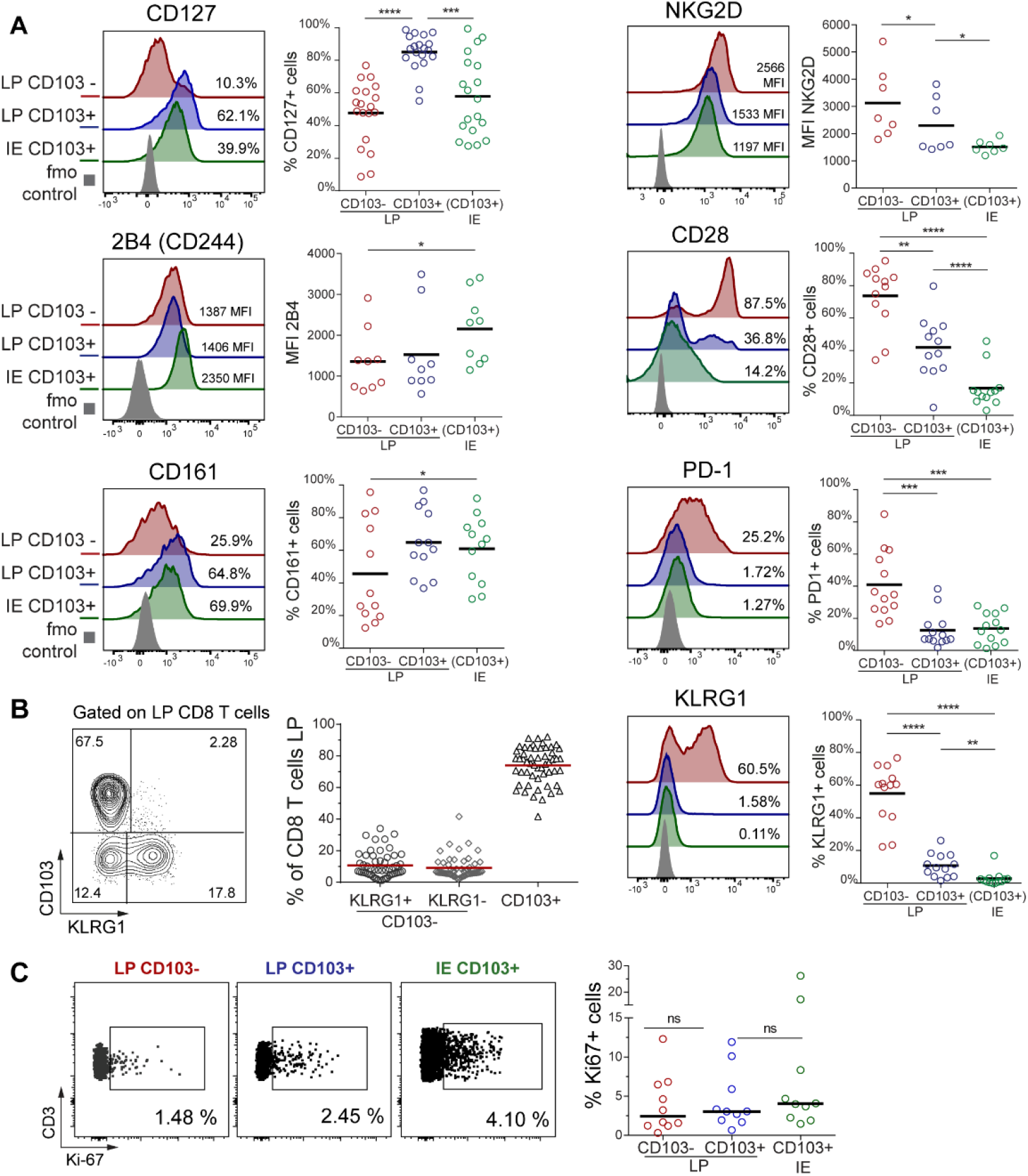
LP and IE CD103+ CD8 T cells are phenotypically distinct from LP CD103-CD8 T cells in normal human small intestine. **(A)** Percentage of positive cells or median fluorescence intensity (MFI) values for various markers on LP CD103-, LP CD103+, and IE CD103+ CD8 T cells in histologically normal small intestine. Representative histograms for all markers are given with color-codes (left): LP CD103-(red), LP CD103+ (blue), IE CD103+ (green) CD8 T cells, fmo control (grey). Black bars indicate median values. (**B**) Representative contour-plot (left) and fraction of LP CD103+ and CD103-CD8 T cells expressing KLRG1 (right, n=54). Red bars indicate mean value. (**C**) Representative dot plot (left) and percentage of LP CD103+, LP CD103-, and IE CD103+ CD8 T cells expressing Ki67 (right graph, n=10). Black bars indicate median values. Statistical analysis was performed using one-way ANOVA for repeated measures with Tukey’s multiple comparisons test. ns, not significant; *,p≤0.05; **,p≤0.01; ***,p≤0.001; ****,p≤0.0001.

To summarize, we showed that the majority of LP CD8 T cells and nearly all IE CD8 T cells express a Trm phenotype (CD103+ KLRG1-), whereas a minor fraction in the LP was more similar to PB CD8 Tem cells being CD103-KLRG1+/-.

### CD8 Trm cells persist for at least one year in transplanted small intestine

Next, we wanted to determine whether CD8 T cells expressing a Trm phenotype were maintained over time in the human SI. The *in vivo* replacement kinetics of CD8 T cells in duodenal biopsies obtained by endoscopy was examined at 3, 6 and 52 weeks after Tx (Horneland et al., 2015) (for details see **Figure S5**). Most donors and patients express different human leukocyte antigen (HLA) type I molecules making it possible to accurately distinguish persisting donor cells from newly recruited incoming recipient cells in the graft by flow cytometry (Landsverk et al., 2017) (**Figure 3A**). Only patients without histological or clinical signs of rejection were included (n=32). At 3 and 6 weeks post-Tx the vast majority of LP and IE CD103+ CD8 T cells were still of donor origin **(Figure 3B)**. Of note, on average more than 60% of both LP and IE CD103+ CD8 T cells were still of donor origin one year after Tx, although there was considerable inter-individual variation with some patients showing hardly any replacement at all (**Figure 3B**). Immunophenotyping showed that the persisting donor CD103+ CD8 T cells expressed a Trm phenotype similar to the baseline situation **(Figure S6**). LP CD103-CD8 T cells were more rapidly replaced by recipient CD8 T cells, and at 6 weeks post-Tx only 34% of the cells were of donor origin, however, after one year the LP CD103-CD8 T cell compartment still contained about 15% donor cells (**Figure 3B**). The KLRG1+ and KLRG1-subsets within the LP CD103-CD8 T cell population exhibited a similar replacement rate at all-time points **(Figure 3C)**. The replacement kinetics of LP and IE CD103+ CD8 T cells were strongly correlated (r^2^=0.82, p<0.0001), whereas those of CD103+ and CD103-CD8 T cells were not (**Figure S7**). Analyzing recipient CD8 T cells we found that 60-70% of incoming LP CD8 T cells were CD103-KLRG1+/-at 3 and 6 weeks post-Tx (**Figure 3D-E**). However, one year after Tx more than half of recipient LP CD8 T cells expressed a Trm phenotype (CD103+ KLRG1-), whereas the relative fractions of CD8 T cell phenotypes were stable at all the time points in the native duodenum (**Figure 3D-E**). This suggested that incoming CD103-CD8 T cells gradually differentiated into CD103+ Trm cells, and that the relative distribution of the CD103+ subset within the total recipient CD8 T cell compartment at one year post-Tx became similar to the steady-state situation (**Figure 2B and 3D-E**). Flow-cytometric analysis of biopsies obtained from the adjacent recipient duodenum did not show any donor-derived CD8 T cells, indicating that lateral migration or propagation from any residual lymphoid tissue in the graft was not occurring **(Figure S8)**.

**Figure 3.**
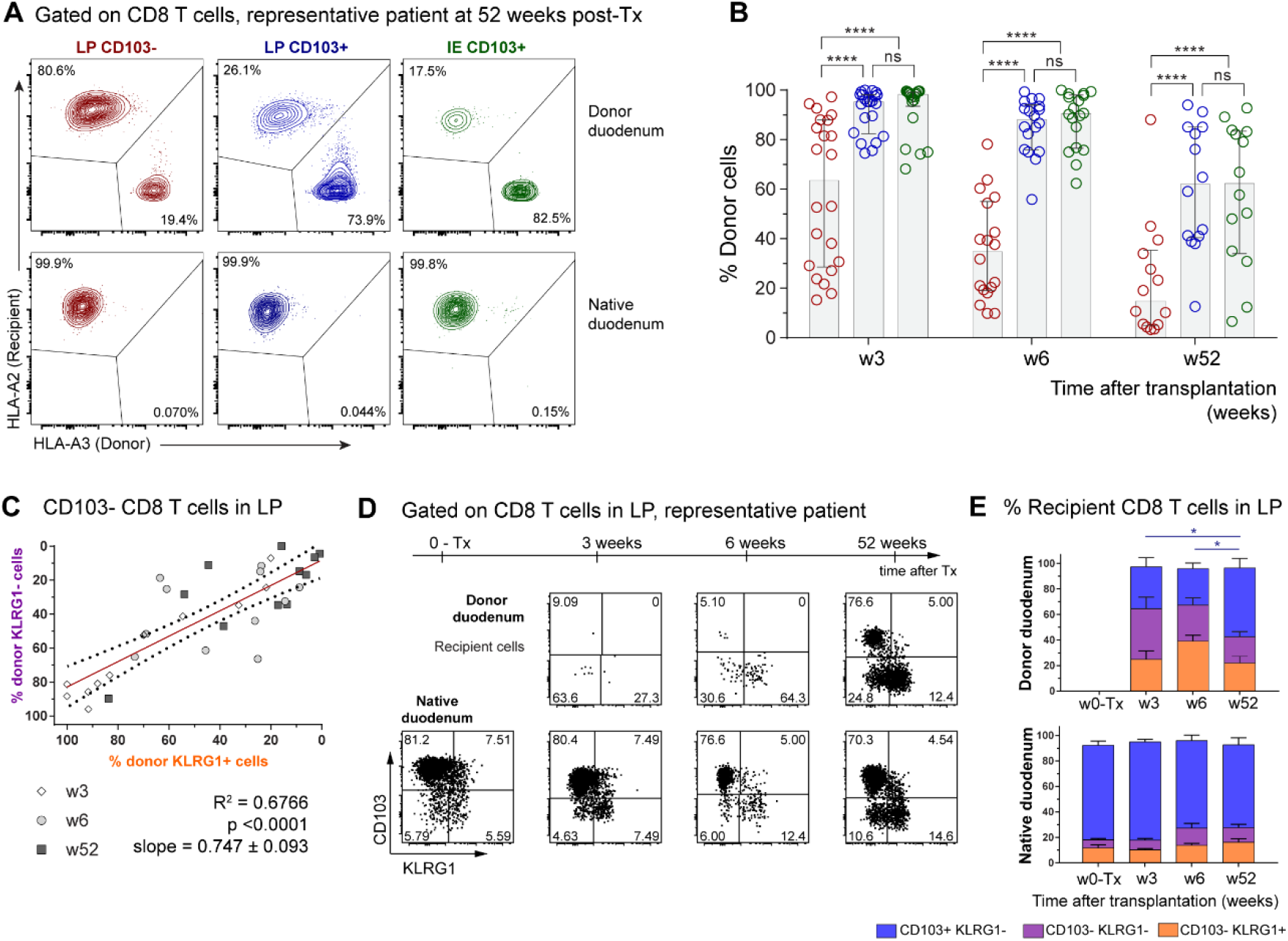
CD103+ CD8 T cells persist for at least one year while CD103-CD8 T cells are dynamically exchanged in transplanted small intestine. (**A**). Representative contour plot showing the percentage of donor and recipient LP CD103+, LP CD103-, and IE CD103+ CD8 T cells in donor duodenum 52 weeks after transplantation. **(B)** Percentage of donor cells in the LP CD103-(left, red), LP CD103+ (center, blue) and IE CD103+ (right, green) CD8 T cells at 3, 6 and 52 weeks after transplantation as determined by HLA class I expression as in (A). Grey columns indicate median values (n=14). (**C**) Correlation of KLRG1+ and KLRG1-subset replacement in the LP CD103-CD8 T cell compartment. Statistics performed using Pearson correlation with two-tailed p-value (95% confidence interval, n=33). (**D**) Distribution of recipient derived CD8 T cells in the LP in different subsets according to the expression of KLRG1 and CD103, in donor (top) and native duodenum (lower), before (0-Tx) and 3, 6 and 52 weeks after transplantation. One representative patient sample is shown. (**E**) Compiled data for recipient CD8 T cell subset representation in native and donor duodenum before (w0-Tx) and after transplantation (w3-52). Statistics was performed using two-way ANOVA with repeated measures across subsets and Tukey’s multiple comparisons test of CD103+ KLRG1-subset with time. ns, not significant; *P≤0.05; ****P≤0.0001.

To evaluate whether the surgical trauma, immunosuppressive treatment and leukocyte chimerism was influencing the absolute numbers of SI CD8 T cells in the transplanted duodenum, we performed immunohistochemical staining of biopsies obtained at different time points from the same patients. Notably, we found that the overall density of CD8 T cells in transplanted duodenum was stable throughout the one-year follow-up period (**Figure S9**). Moreover, flow-cytometric analysis of Tx tissue showed very few Ki67+ cells among the donor LP and IE CD103+ CD8 T cells (**Figure S10A-B**), similar to steady-state levels **(Figure 2C)**, indicating that proliferation was not a major contributing factor to the persistence of donor T cells in transplanted intestine.

Taken together, these results show that SI CD103+ CD8 T cells can survive for at least one year in the tissue and suggest that CD103-CD8 T cells recruited to the SI may differentiate into Trm cells *in situ*.

### Single-cell TCR repertoire analysis supports long-term persistence of CD103+ CD8 T cells in transplanted intestine

To further explore and verify the long-term persistence of SI CD8 Trm cells in Tx, we studied the conservation of the immune repertoire of SI CD103+ CD8 T cells over time at the single-cell level. Specifically, we performed single-cell high throughput TCR sequencing of donor LP CD103+ CD8 T cells sorted from the grafted duodenum before and one year after Tx. To determine if the degree of chimerism might affect the TCR repertoire, we included samples from one patient exhibiting high T-cell replacement one year post-Tx (Ptx#1; 5% donor CD8 T cells) and one patient with low replacement (Ptx#2; 70% donor CD8 T cells, for sorting gating strategy see **Figure S11**). As a control we also performed single-cell TCR sequencing of LP CD103+ CD8 T cells from biopsies of the native (i.e. autologous, non-transplanted) duodenum of one patient (Ptx#2).

First, we investigated how many cells with the same clonotype (defined by identical nucleotide TCRα+β chains) were present in samples from the same individual taken before and one year post-Tx. Strikingly, despite the limited sampling, we detected overlapping clonotypes in biopsies obtained one year apart both in transplanted and non-transplanted tissue. We found that 12% of the clonotypes identified at baseline were present one-year post-Tx in patient Ptx#1, whereas a 50% and 37% overlap was detected in patient Ptx#2 (donor and native duodenum, respectively) (**Figure 4A**). The frequency distribution of the TCRα-β clonotypes at both time points was similar in all the paired samples, and the first 10 clones with higher clonal size (8 out of 21 clonotypes and 10 out of 35 clonotypes on average, at w0 and w52, respectively) represented around half of the repertoire in all the cases, indicating significant clonal expansion **(Figure 4B**). Most of the unique TCRα-β sequences were derived from relatively small clones (expressed only by one cell), whereas a higher proportion of expanded clones was found among those shared at both time points (**Figure 4C**). In agreement with other reports (Han et al., 2014), we found that TCR-sequencing efficiency was consistently lower for TCRα. In order to analyze a larger number of cells we therefore calculated the clonal overlap using only the TCRβ chain sequence. In all paired samples the percentage of overlapping TCRβ clonotypes was comparable to that calculated for TCRα-β clonotypes (**Figure 4D**). Furthermore, TCRβ analysis showed that the shared clones were more expanded both at baseline and at 52 weeks compared to the unique clones (**Figure 4E**). The degree of clonal overlap and the frequency of expanded clones were similar in the grafted and the native duodenum from the same patient (Ptx#2), suggesting that chimerism in the transplanted SI did not trigger an excessive allo-driven clonal expansion. This was substantiated by the finding that CD8 T cells showed low expression of the proliferation marker Ki67 (**Figure S10A-B).**

**Figure 4.**
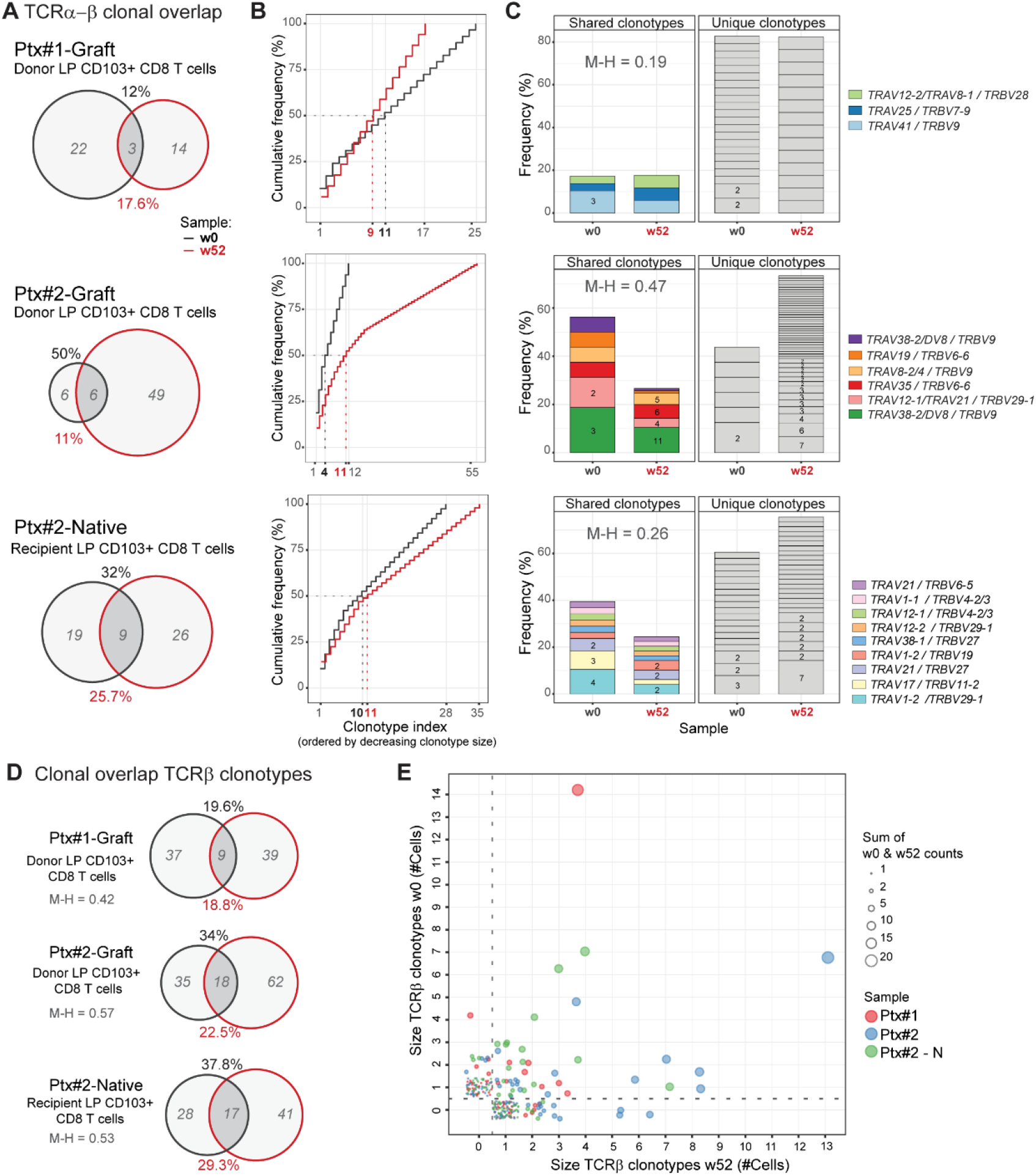
Persisting clonotypes of CD103+ CD8 Trm cells are detected in SI samples one-year after Tx. **(A)** Clonotype overlap analysis by single-cell TCR-sequencing of donor CD103+ CD8 Trm cells derived from donor duodenum at baseline (w0, black) and one-year after transplantation (w52, red) (Ptx#1- and Ptx#2-Donor) and of native CD103+ CD8 Trms from native duodenum (Ptx#2-Native). In italics (gray), number of unique clonotypes. Clonotype overlap was calculated as follows: %overlap(X,Y)=(|X ∩Y|)/((|X| or |Y|))*100, where X represents the total clonotypes at w0 (% in black) and Y represents the total clonotypes at w52 (% in red). **(B)** Frequency distribution of LP CD103+ CD8 Trm clonotypes at baseline (w0, black line) and 1-year post-Tx (w52, red line). Clonotype index is ordered by decreasing clonotype size. **(C)** Clonal dominance and preferential *TRAV/TRBV* pairing. Each slice of the column represents a different *TRAV/TRBV* pair and the number of cells expressing them is shown (for >1 cell). Unique and overlapping *TRAV/TRBV* clones are shown separately.(**D**) Clonotype overlap analysis of total TCR-B clonotypes in the same samples as described in (**A**). (**E**) Correlation of the TCR-B clonotypes found at baseline (w0) and one year after transplantation (w52). The samples are color-coded and clonal size is given. M-H, Morisita-Horn index.

Together, these results confirmed that persisting CD8 Trm cells survive for at least one year in transplanted duodenum, and showed that relatively few expanded clonotypes constitute a considerable fraction of the CD8 Trm repertoire.

### LP and IE CD103+ Trm cells present similar immune repertoire

The replacement kinetics and phenotype of CD8 T cells in transplanted SI **(Figure 3B and Figure S6**) suggested that both IE and LP CD103+ CD8 T cells are Trm cells, whereas CD103-KLRG1+/-CD8 T cells were more dynamic subsets that have the potential to differentiate into CD103+ CD8 T cells (**Fig 3D-E**). To further interrogate the relationship among the SI CD8 T cell subsets, we sorted LP and IE CD103+ CD8 T cells as well as KLRG1+ and KLRG1-LP CD103-subsets from five SI samples. RNA was isolated and TCRα and TCRβ genes were amplified by semi-nested PCR including three cDNA replicates for each TCR chain (**Figure S12A**). Correlation of the read counts for the clonotypes found within the three molecular replicates was high **(Figure S12C)**, indicating reproducibility of the TCR repertoire sequencing (Greiff et al., 2014). In order to determine whether different SI CD8 T cell subsets shared a common clonal origin we calculated the percentage of overlapping clones and the Morisita-Horn similarity index for all the pairwise comparisons of both TCRα and TCRβ clonotypes. We found that the repertoire of LP and IE CD103+ CD8 T cells was very similar with a clonal overlap of 35.6% and 37.5% and Morisita-Horn indexes of 0.66 and 0.71, respectively (**Figure 5A-B and Figure S13)**. Both CD103-CD8 T cell subsets had a relatively low degree of similarity with the CD103+ subsets. However, notably, the Morisita-Horn index was 10-fold higher comparing LP and IE CD103+ CD8 T cells with the KLRG1-subset than with the KLRG1+ subset (**Figure 5B** and **Figure S13)**. Similar values of clonal overlap and Morisita-Horn indexes were obtained when the CDR3 amino acid sequence was used to define the clonotypes (**Figure S14**).

**Figure 5.**
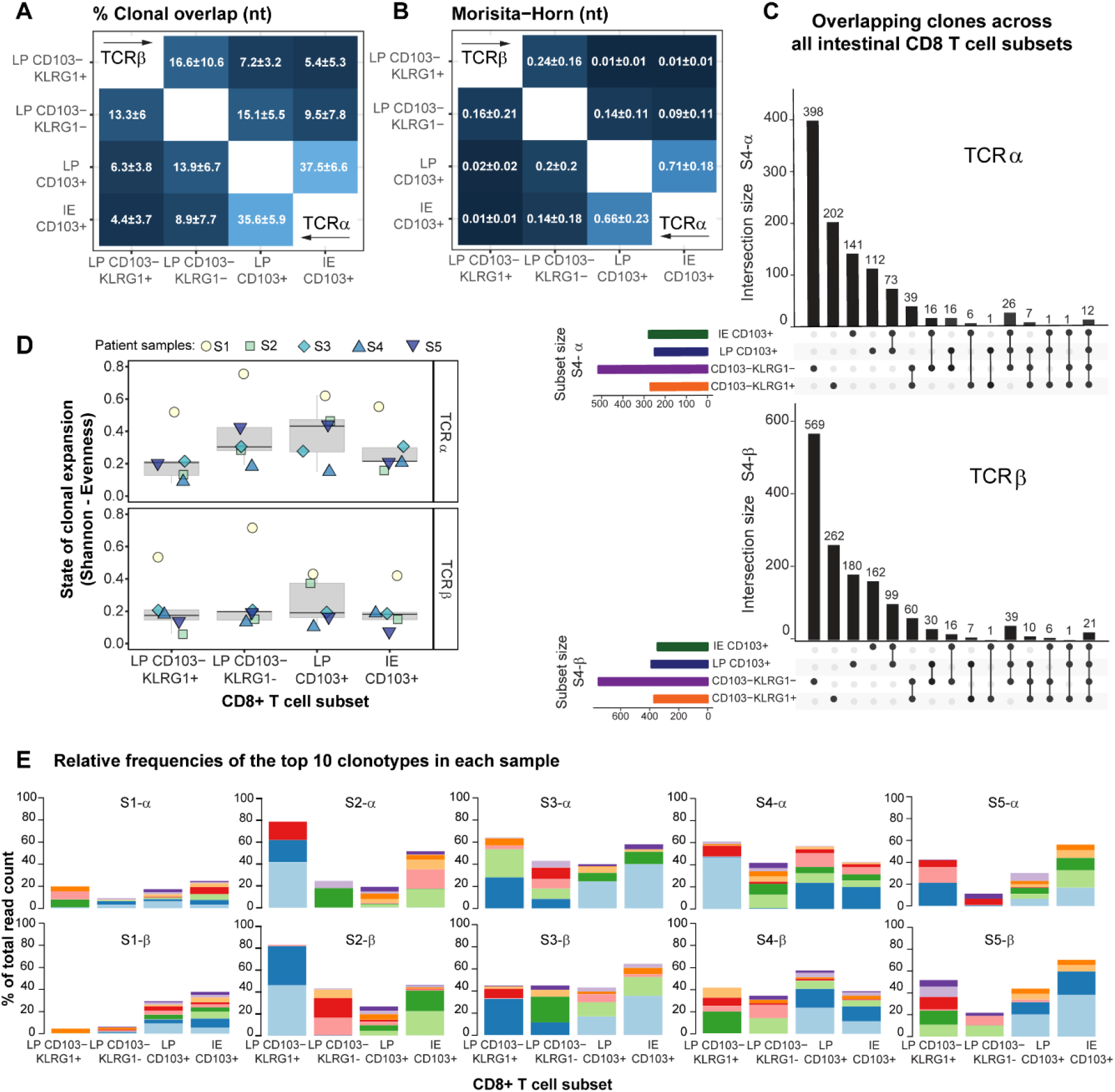
High clonal overlap between IE and LP CD103+ CD8 T cells. **(A)** Mean percentage of clonal overlap and **(B)** Morisita-Horn similarity indexes applied to all the pairwise combinations of CD8 T cell subsets for TCRα (lower corner) and TCRβ clonotypes (upper corner) derived from normal SI (n=5). The Morisita-Horn index ranges from 0 (no similarity) to 1 (identical). **(C)** Overlapping clones among the different intestinal CD8 T cell subsets for TCRα and TCRβ in one representative sample (n=5, **Figure S14**). Intersections are represented below the x axis (black circles), the number of overlapping clonotypes is represented on the histogram and the total amount of clonotypes per subset as horizontal bars **(D)**. Evenness index is calculated for the TCRα and TCRβ repertoire of each CD8 T cell subset per sample. Values range from 0 (monoclonal distribution) to 1 (uniform distribution). **(E)** Size of the read counts for the 10 most expanded clonotypes relative to the total reads in each subset. Shared clones are represented with the same color within the same sample and TR chain.

To better understand the extent of clonal overlap within the SI CD8 T cell population we measured overlapping clones across all the SI CD8 T cell subsets. Interestingly, we found that some clones were shared among three or even all four CD8 T cell subsets (representative sample in **Figure 5C** and all the samples in **Figure S15**), suggesting that the same progenitor cell can give rise to all memory CD8 T cell subsets. We next assessed the clonal diversity for each subset by calculating the Shannon-Evenness index, which describes the extent to which a distribution of clonotypes is distanced from the uniform distribution, with values ranging from 0 (least diverse, clonal dominance) to 1 (highly diverse, all clones have the same frequency), (Greiff et al., 2015b). We found that all four SI CD8 T cell subsets displayed relatively low evenness values (**Figure 5D**), indicating a polarized repertoire with prevalence of certain clones. **Figure 5E** shows the abundance of the 10 most expanded TCRα and TCRβ clonotypes in each sample. Notably, a few dominant clones constituted a large fraction of the total repertoire in all of the subsets (up to 40% of the total reads in some cases). In line with their similarity values (**Figure 5B**), LP and IE CD103+ T cells showed an analogous distribution of 10-top ranked clones in all 5 donors (**Figure 5E**).

To summarize, these data show that LP and IE CD103+ Trm cells have a very similar TCR repertoire, in both cases this repertoire is biased towards a few expanded clones, and it is more closely related to the LP CD103-CD8 T cell subset lacking KLRG1 expression.

### Activated SI CD8 T cell subsets produce different levels of cytokines and cytotoxic mediators

CD8 T cells exert their effector functions through activation-induced cytotoxicity involving targeted secretion of perforin and granzyme-B and cytokine secretion. In the absence of stimulation significantly more CD103-CD8 T cells expressed granzyme-B and perforin compared to LP and IE CD103+ T cells (**Figure 6A**). However, after activation with anti-CD3/CD28 beads, both CD103-CD8 T cells and LP CD103+ CD8 T cells significantly increased their expression of granzyme-B and perforin, although the level of perforin was still higher in the CD103-CD8 T cell subset (**Figure 6B)**. Within the CD103-population, KLRG1+ cells expressed higher levels of perforin after stimulation compared to KLRG1-cells (**Figure S16A-B**). Interestingly, induction of cytolytic molecules was hardly detectable in IE CD8 T cells, suggesting that TCR stimulation is not sufficient to activate their cytolytic function (**Figure 6B**).

**Figure 6.**
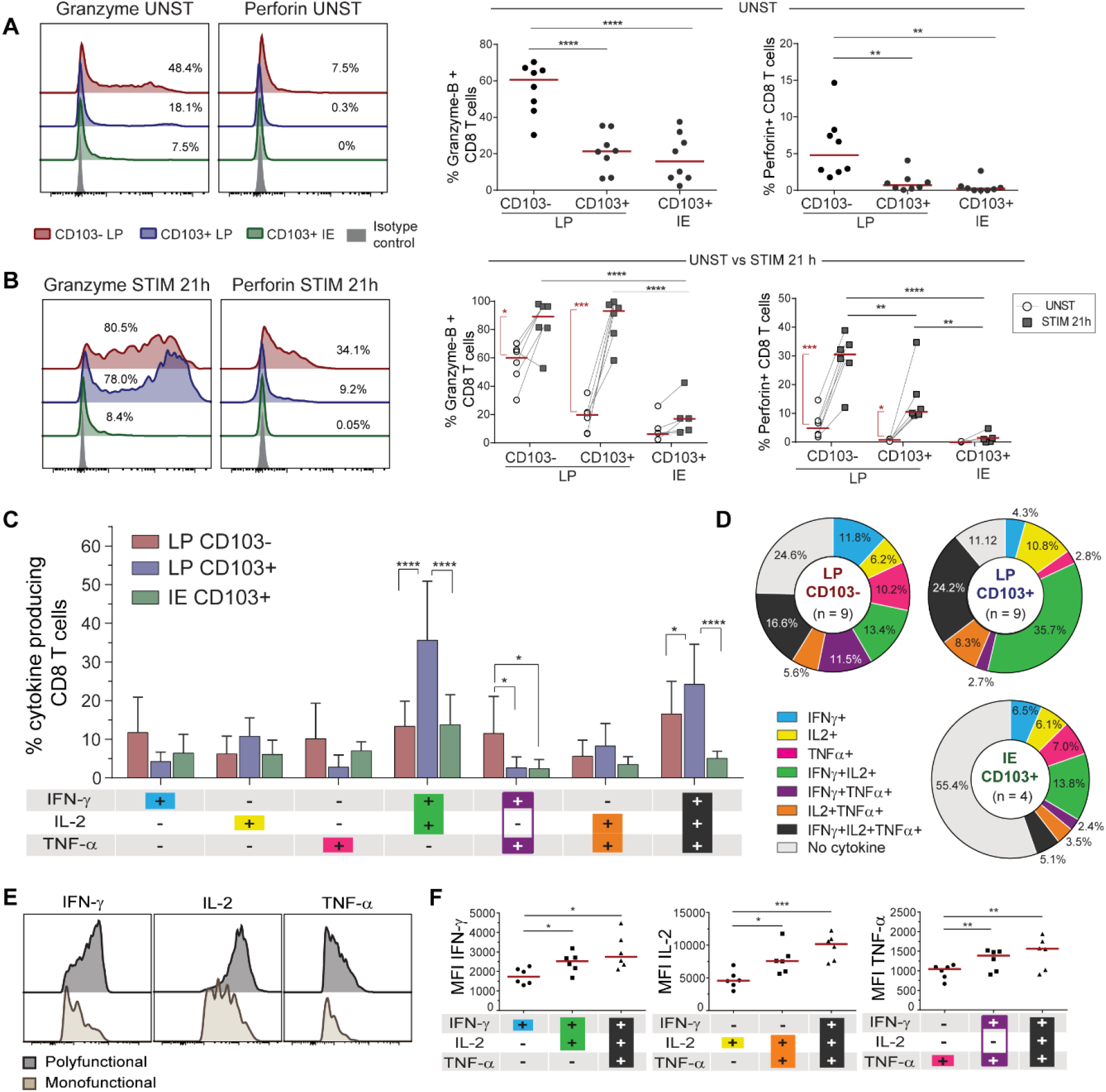
SI CD8 T cell subsets produce different levels of cytokines and cytotoxic molecules in response to activation. (**A, B**) Representative flow-cytometric histogram (left) and compiled data (right, n=8) for the intracellular expression of granzyme-B and perforin in CD8 T cell subsets (color-coded) and isotype control (grey) without (A) and after (**B**) stimulation with anti-CD3/CD28 beads for 21h (n=6). (**C**) Flow-cytometric analysis of PMA/ionomycin induced cytokine production by LP CD103-, LP CD103+ and IE CD103+ CD8 T cells. The mean percentages of cytokine producing cells with SD are given by bars. (**D**) Relative representation of specific cytokine production profiles for LP CD103- and LP CD103+ and IE CD103+ CD8 T cells are represented on pie charts with color code as beneath x-axis in (**C**). Mean values of indicated experiments. (**E**) Representative histograms and (**F**) compiled median fluorescence intensity (MFI) values for single-producing, dual- and triple-cytokine producing CD8 T cells (n=6). Statistical analysis was performed using one-way (A, F) and two-way (B, C) ANOVA-RM with Tukey’s multiple comparisons test. For CD3/CD28 stimulation in (B) Students T-test was applied to compare unstimulated and stimulated cells (red vertical lines and asterisks). *P≤0.05, **P≤0.01, ***P≤0.001, ****, p≤0.0001. Red horizontal lines on graphs represent median values.

To investigate their cytokine production profiles, SI CD8 T cell subsets were subjected to short-term stimulation with phorbol-12-myristate-13-acetate (PMA) and ionomycin, followed by intracellular flow-cytometric cytokine detection. Most (88%) LP CD103+ CD8 T cells produced one or more of the cytokines tested, whereas 75% of the LP CD103-CD8 T cells and 45% of the IE CD8 T cells were positive (**Figure 6C-D**). LP CD103+ CD8 T cells had a significant higher proportion of dual IFN-γ+IL-2+ cells and triple-producing (IFN-γ+IL-2+TNF-α+) cells compared to the other subsets (**Figure 6C**). Moreover, triple-producing CD8 T cells produced significantly higher levels of the individual cytokines compared to single cytokine-producing cells (**Figure 6E-F**). Within the CD103-CD8 T cell population we found that KLRG1-cells contained significantly higher numbers of triple-producing cells than their KLRG1+ counterpart (**Figure S16C**).

Taken together, these data showed that CD8 Trm cells in LP are more potent cytokine-producing cells than CD103-CD8 T cells. All LP CD8 T cell subsets efficiently produced cytotoxic molecules after activation, whereas IE CD8 Trm cells produced significantly less cytokines and cytotoxic molecules.

## Discussion

Here we show that the human small intestine contains several phenotypically, functionally and clonally distinct CD8 T cell subsets. Virtually all IE CD8 T cells and the majority of LP CD8 T cells expressed a Trm phenotype and were persistent cells (>1 year) with very similar immune repertoire. When activated, all subsets contained cytokine-producing cells, with LP CD8 Trm cells being particularly efficient. The two minor CD103-CD8 T cell subsets were phenotypically more similar to PB CD8 T cells, presented an immune repertoire more different from the Trm cells, and produced higher levels of granzyme-B and perforin.

Experiments to determine the longevity of lymphocytes are not readily performed in human tissues (Landsverk et al., 2017); however, by studying the replacement kinetics in a transplantation setting we were able to determine the persistence of CD8 T cells in grafted human duodenum. By multi-parameter flow-cytometric analysis of CD8 T cells derived from normal (non-transplanted) SI we found that LP CD8 T cells could be divided into three distinct subsets depending on their expression of CD103 and KLRG1. Most of CD8 T cells displayed a Trm phenotype (CD103+ KLRG1-). In contrast, the two minor CD103-subsets in LP being either KLRG1- or KLRG1+ were more similar to the CD8 T cells in peripheral blood, presenting higher levels of CD28, PD1, NKG2D and lower CD127 and CD161 expression. Following the turnover kinetics of the subsets in transplanted duodenum we found in half of the patients that more than 60% of both IE and LP CD103+CD8 T cells were of donor origin one year after transplantation with no reduction in absolute numbers of CD8 T cells. The finding of persistent donor CD8 T cells was substantiated by single-cell TCR sequencing showing a significant clonal overlap between donor LP CD103+ CD8 T cells obtained before and one year post-transplantation. Donor CD103-CD8 T cells, in contrast, were more rapidly replaced by recipient cells. However, interestingly, a small fraction (15%) of the CD103-subsets were still of donor origin one year after transplantation.

Zuber et al. (Zuber et al., 2016) reported recently that donor CD8 T cells were present in some non-rejected intestinal transplants for more than 600 days. These patients received intestinal transplantation alone or as part of multivisceral transplantation, which in both situations include gut-associated lymphoid tissue (GALT, e.g. Peyers patches) and mesenteric lymph nodes. Long-term mixed chimerism is observed in the blood of these patients (Fu et al., 2018), suggesting that organized lymphatic tissue in the graft continuously expand and release donor T cells that “home” to the intestinal mucosa. In our study, however, only the proximal part of the duodenum was transplanted without GALT or lymph nodes, excluding the possibility of continuous replenishment of donor cells after transplantation.

Our findings show that CD8 T clones have the capacity to survive for at least a year and most probably many years in the human intestinal mucosa, thus demonstrating that *bona fide* Trm cells exist in the human SI. In the steady state, Trm can be generated from circulating memory precursors or proliferate locally in response to cognate antigens (Beura et al., 2018a). The proliferation rate, as measured by Ki67 staining, was consistently low in the transplanted and non-transplanted SI. This is in agreement with previous reports (Thome et al., 2014), suggesting that the persistence of CD8 T cells was maintained by long-lived cells rather than local proliferation. The finding that there was no massive expansion of selected clones one-year post-transplantation compared with the situation at baseline and with the native duodenum supports this scenario. Most of the persistent CD8 T cells expressed the classical Trm marker CD103 (αE integrin), that mediates homotypic interactions with epithelial cells via E-cadherin (Schon et al., 1999). However, a small population of CD103-CD8 T cells was also maintained for one year in the transplanted duodenum, suggesting that CD103 is not an obligate marker for all Trm cells, as has been suggested also for intestinal CD8 T cells in mice (Bergsbaken and Bevan, 2015).

Translated to the steady state situation it is likely that the turnover rate of CD8 T cells observed in transplanted SI underestimates their physiological longevity for several reasons. First, all patients received immunosuppressive treatment including anti-thymoglobulin that rapidly depletes T cells in blood (Horneland et al., 2015). It is likely that this treatment also affects the T cell numbers in the periphery. Second, solid organ transplantation surgery causes ischemia and reperfusion injury of the graft that increases the turnover of leukocytes (Eguiluz-Gracia et al., 2016); and third, our previous studies have shown that dendritic cells and some of the macrophage subsets are rapidly replaced in the transplanted duodenum (Bujko et al., 2018; Richter et al., 2018), thereby increasing the risk of local allo-reactivity. As shown by Zuber et al (Zuber et al., 2016) the turnover rate of T cells increases in intestinal tissue with graft rejection. Although patients with histological signs of rejection were excluded from the study, all included patients displayed T cell recipient chimerism in the graft, in which episodes of low grade rejection cannot be excluded. Altogether, our results, derived from both transplanted and normal SI combined, strongly suggest that virtually all IE CD8 T cells and the majority of LP CD8 T cells under homeostatic conditions are long-lived Trm cells with low proliferative activity (<5% Ki67+ cells).

Most studies of SI CD8 T cells in humans have focused on the intraepithelial compartment, and there is less knowledge about the CD8 T cell population in LP. Here we show that IE and LP CD103+ CD8 T cells present a very similar immune repertoire, indicating a common origin. When stimulated with PMA/ionomycin about 40% of IE CD8 T cells produced cytokines (IFN-γ, IL-2 or TNF-α), whereas nearly all LP CD8 Trm cells were cytokine-producing, and more than a third of these cells produced IFN-γ, IL-2 and TNF-α simultaneously (polyfunctional). This enhanced ability of Trm to produce robust responses mediated by polyfunctional cytokine production have been associated with the role of Trm as sentinel cells in different barrier tissues, such as skin (Watanabe et al., 2015). When activated with anti-CD3/CD28 beads LP CD8 Trm cells produced high amounts of granzyme-B and significant levels of perforin, whereas IE CD8 T cells expressed less of these cytotoxic mediators. This finding may to some extent be explained by the fact that IE CD8 T cells expressed only low levels of CD28. This suggests that intestinal CD8 Trm cells have a unique ability to convey strong and long-term protective immunity against pathogens. Although IE CD8 T cells were less responsive to the stimuli we used in this study, their TCR repertoire was similar to LP CD103+ CD8 T cells, indicating that they had been activated under similar antigenic conditions. Moreover, in the transplanted intestine we found that the turnover kinetics of IE and LP CD103+ CD8 T cells were strongly correlated (Figure **S6**), together suggesting that IE and LP CD8 Trm cells were interrelated populations rather than operating independently in different compartments. Mouse models have shown that intestinal T cells are very motile and can move back and forth between the LP and the epithelium (Hu et al., 2018) and this may also be the case for CD8 T cells in the human gut. In addition, epithelial cells are in continuous communication with IE CD8 T cells, and it is therefore plausible that epithelial cell signaling under steady state conditions might increase the threshold of TCR-mediated response as a mechanism to maintain the integrity of the epithelial barrier (Jabri and Ebert, 2007).

Approximately 20% of SI CD8 T cells in LP did not express CD103 under steady state conditions. They were phenotypically more similar to PB CD8 T cells than CD8 Trm cells and the immune repertoire of CD103-CD8 T cells showed low clonal overlap with their Trm counterparts. In transplanted intestine incoming recipient CD8 T cells were mainly CD103-at 3 and 6 weeks post-transplantation, which is compatible with the notion that CD103-CD8 T cells in the steady state are mostly recently recruited cells. However, one-year post-transplantation most recipient CD8 T cells presented a Trm phenotype, indicating that CD103-CD8 T cells differentiated into CD8 Trm cells *in situ*. Comparing the TCR repertoire, CD103-KLRG1-CD8 T cells were more similar to CD8 Trm cells than CD103-KLRG1+ cells, which accords with studies in mice showing that KLRG1-CD8 T cells are Trm precursors (Mackay et al., 2013; Sheridan et al., 2014). The KLRG1-subset was also more similar to Trm cells with regard to production of cytokines and cytotoxic molecules. Notwithstanding, it has been shown that TGF-β, abundantly present in the intestinal microenvironment, induces downregulation of KLRG1 (Schwartzkopff et al., 2015). Thus, it is possible that also KLRG1+ T cells can differentiate into Trm cells (Herndler-Brandstetter et al., 2018).

Interestingly, the immune repertoire in both CD103+ and CD103-subsets was polarized towards few dominating expanded clones. Assuming that CD103-CD8 T cells were recently recruited from the circulation this finding suggests that the clonal expansion had occurred before disseminating into the tissue, most likely in GALT and mesenteric lymph nodes. In agreement with this concept, biopsies obtained from different sites (several cm apart) and at different time points in transplanted patients showed expansion of many of the same clones, suggesting that T cell clones were evenly distributed along the duodenal mucosa and less dependent on local proliferation. Thus, the maintenance of a polarized immune repertoire (likely specific to recurrent pathogens) conserved both longitudinally in the tissue and over time, represent an optimized strategy of protection.

There is increasing evidence that Trm cells play an important role in mediating protective responses and maintaining long-term immunity against a broad variety of infectious diseases (Muruganandah et al., 2018). These studies are mainly performed in mouse models and studies investigating the existence and function of Trm cells in human tissue are still few. Here we provide evidence that the majority of SI CD8 T cells in the steady state are long-lived cells with very highly potent protective capabilities. These data indicate that Trm cells in the intestinal mucosa are attractive targets to design effective oral vaccines. Further, the knowledge that long-lived T cells exist in the gut should give incentive for the implementation of new therapeutic strategies in diseases where pathogenic T cells play an important role, such as inflammatory bowel disease, celiac disease and gut graft versus host disease.

## Materials and Methods

### Human biological material

Small intestinal samples were either obtained during pancreatic cancer surgery (Whipple procedure, n = 35; mean age 63yr; range 40-81yr; 16 female), or from donors and/or patients during pancreas-duodenum transplantation (donors: n = 52; mean age 31yr; range 5-55yr; 24 female; patients: n = 36; mean age 41yr; range 25-60yr; 14 female) as described previously (Landsverk et al., 2017; Richter et al., 2018). Endoscopic biopsies from donor and patient duodenum were obtained 3, 6 and 52 weeks after transplantation. All transplanted patients received a standard immunosuppressive regimen (Horneland et al., 2015). Cancer patients receiving neoadjuvant chemotherapy and transplanted patients showing evidence of rejection were excluded from the study. All tissue specimens were evaluated blindly by experienced pathologists, and only material with normal histology was included (Ruiz et al., 2010). Blood samples were collected at the time of the surgery and buffy coats from healthy donors (Oslo University Hospital).

Intestinal resections were opened longitudinally and rinsed with PBS, and mucosa was dissected in strips off the submucosa. For microscopy, small mucosal pieces were fixed in 4% formalin and embedded in paraffin according to standard protocols. Intestinal mucosa was washed 3 times in PBS containing 2mM EDTA and 1% FCS at 37°C with shaking for 20 minutes and filtered through nylon 100-µm mesh to remove epithelial cells. Epithelial fractions in each washing step were pooled and filtered through 100-µm cell strainers (BD, Falcon). After three sequential washes with EDTA buffer, the epithelial layer was completely removed from the tissue and the basal membrane remained intact as it is shown in **Figure S1A**. Epithelial cells in the EDTA fraction were depleted by incubation with anti-human epithelial antigen antibody (clone Ber-EP4, Dako) followed by anti-mouse IgG dynabeads (ThermoFisher) according to the manufacture’s protocol. A death cell removal kit (Miltenyi), was applied to the depleted IE cells before cell sorting and functional assays. LP was minced and digested in complete RPMI medium (supplemented with 10% FCS, 1% Pen/Strep) containing 0.25 mg/mL Liberase TL and 20 U/mL DNase I (both from Roche), stirring at 37°C for 1h. Digested tissue was filtered twice through 100-µm cell strainers and washed tree times in PBS. Purity of both IE and LP fractions was checked by flow-cytometry. We found <5% BerEP4+ epithelial cells in LP and no expression of B cell / plasma cell markers (CD19, CD27) in the IE (**Figure S1B-C**), confirming that the degree of cross-contamination between these fractions was very low. Intestinal biopsies from transplanted patients were processed following the same protocol. PBMCs were isolated by Ficoll-based density gradient centrifugation (Lymphoprep™, Axis-Shield). All participants gave their written informed consent. The study was approved by the Regional Committee for Medical Research Ethics in Southeast Norway and complies with the Declaration of Helsinki.

### Microscopy

Analysis of chimerism was performed as described previously (Landsverk et al., 2017). Briefly, formalin-fixed 4-μm sections were washed sequentially in xylene, ethanol, and PBS. Heat-induced epitope retrieval was performed by boiling sections for 20min in Dako buffer. Sections were incubated with CEP X SpectrumOrange/Y SpectrumGreen DNA Probes (Abbott Molecular Inc.) for 12h at 37°C before immunostaining according to standard protocol with anti-CD3 (Polyclonal; Dako), anti-CD8 (clone 4B11, Novocastra) and secondary antibodies targeting rabbit IgG or mouse IgG2b conjugated to Alexa Fluor 647 and 555, respectively. Laser scanning confocal microscopy was performed on an Olympus FV1000 (BX61WI) system. Image z stacks were acquired at 1-μm intervals and combined using the Z project max intensity function in Image J (National Institutes of Health). All microscopy images were assembled in Photoshop and Illustrator CC (Adobe).

CD8 immunoenzymatic staining was performed on formalin-fixed 4-μm sections, dewaxed in xylene and rehydrated in ethanol, and prepared with Vulcan Fast red kit (Biocare Medical) following standard protocols. In brief, heat-induced antigen retrieval was performed in Tris/EDTA pH9 buffer, followed by staining with primary antibody (CD8 clone 4B11, Novocastra), secondary anti-mouse AP-conjugated antibody and incubation with substrate (Fast red chromogen, Biocare Medical). Slides were counterstained with hematoxylin and excess of dye was removed by bluing in ammoniac water. Tissue sections were scanned using Pannoramic Midi slide scanner (3DHISTECH) and counts generated with QuPath software (Bankhead et al., 2017).

### Flow cytometry and cell sorting

Immunophenotyping was performed on single cell suspensions of LP and IE fractions and PBMCs using different multicolor combinations of directly conjugated monoclonal antibodies (**Table S1**). To assess the expression of L-Selectin on digested tissue, cells were rested for 12h at 37°C before the immunostaining. Replacement of donor cells in duodenal biopsies of HLA mismatched transplanted patients was assessed using different HLA type I allotype-specific antibodies targeting donor- and/or recipient-derived cells, and stroma cells were used as a control of specific staining. Dead cells were excluded based on propidium iodide staining (Molecular Probes, Life Technologies). For analysis of cytokine production, LP and IE cell suspensions were stimulated for 4h with control complete medium (RPMI supplemented with 10% FCS, 1% Pen/Strep) or phorbol-12-myristate-13-acetate PMA (1.5 ng/mL) and ionomycin (1µg/mL; both from Sigma-Aldrich) in the presence of monensin (Golgi Stop, BD Biosciences) added after 1h of stimulation to allow intracellular accumulation of cytokines. Cells were stained using the BD Cytofix/Cytoperm kit (BD Biosciences) according to the manufacturer’s instructions, and stained with antibodies against TNFα, IFNγ, or IL-2 (**Table S1**). For detection of cytotoxic granules, LP and IE cells were activated for 21h with anti-CD3/CD28 beads (Dynabeads, ThermoFisher) or control complete medium. For detection of intranuclear Ki67 expression the FoxP3/transcription factor staining buffer set was used according to the manufacturer’s instructions. eFluor-450 or eFluor-780 fixable viability dyes (eBioscience) were used prior any intracellular/intranuclear staining procedure. All samples were acquired on LSR Fortessa flow cytometer (BD Biosciences), using FACSDiva software (BD Biosciences). Single stained controls were prepared for compensation (UltraComp eBeads(tm), eBioscience), and gates were adjusted by comparison with FMO controls or matched isotype controls. Flow cytometry data were analyzed using FlowJo 10.4.2 (Tree Star). For **Figure 1C**, the expression of 16 phenotypic markers was analyzed at the single cell-level and compared for CD8 T cells in PB, LP and IE (n=5) using the merge and calculation functions of Infinicyt software (Cytognos), as described in detail elsewhere (Pedreira et al., 2013). The population within the CD8 T cell gate was down-sampled for each compartment and exported to a new file, and then concatenated and subjected to t-SNE analysis using the plugin integrated in FlowJo 10.4.2 (See **Figure S3** for more details). FACS sorting was performed on Aria II Cell Sorter (BD Biosciences). A TCRγd antibody was used to exclude these cells during sorting (see gating strategy in **Figure S11**). All experiments were performed at the Flow Cytometry Core Facility, Oslo University Hospital.

### *Single-cell TCR*α-β *sequencing*

Cell suspensions from two donors (Ptx#1 and Ptx#2) were prepared at the time of transplantation and kept frozen in liquid nitrogen. One-year post-transplantation, biopsies from the same patients (donor and recipient tissue) were collected, and single cell suspensions were prepared and processed together with the thawed baseline samples from the same patients. Donor CD103+ KLRG1-CD8 T cells from LP of transplanted tissue were identified following the gating strategy in **FigureS11** and single cells were sorted into 96-well plates (BioRad) containing 5 µL capture buffer (20 mM Tris-HCl pH8, 1% NP-40, 1 U/µL RNase Inhibitor). The plates were spun-down and snap frozen immediately after sorting and stored at −80°C until cDNA synthesis. Paired TCRα and TCRβ sequences were obtained after three nested PCR with multiplexed primers covering all TCRα and TCRβ V genes, as described before (Risnes et al., 2018). In brief, cDNA plates were stored at −20°C, and each of the three nested PCR steps was carried out in a total volume of 10 µL using 1 µL cDNA/PCR template and KAPA HiFi HotStart ReadyMix (Kapa Biosystems). In the last PCR reaction, *TRAC* and *TRBC* barcoding primers were added together with Illumina PairedEnd primers. Cycling conditions, concentrations and primer sequences for all three PCR reactions can be found in (Risnes et al., 2018), and original protocol in (Han et al., 2014). Products in each well were combined, purified and sequenced using the Illumina paired-end 250 bp MiSeq platform at the Norwegian Sequencing Centre (NSC), at Oslo University Hospital. The resulting paired-end sequencing reads were processed in a multistep pipeline using selected steps of the pRESTO toolkit (Vander Heiden et al., 2014) according to (Risnes et al., 2018). In short, high-purity reads (average Phred score >30) were deconvoluted using barcode identifiers and collapsed, and only the top three for each well were retained for further analysis. For identification of V, D, and J genes and the CDR3 junctions the International ImMunoGeneTics Information System (IMGT)/HighV-QUEST online tool (Alamyar et al., 2012) was used. The results were then filtered and collapsed. Paired TCR sequences were grouped in clonotypes, defined by identical V and J-gene (subgroup level) together with identical CDR3 nucleotide sequence for both TCRα and TCRβ when that information was available. Only valid singleton cells containing no more than 3 chains (dual T cell receptor α [TCRα] and T cell receptor β [TCRβ]) with 100 or more reads were considered for downstream analysis.

### Bulk TCR sequencing

5000 cells of each subset of CD8 T cells (CD103-KLRG1+, CD103-KLRG1- and CD103+ from LP and CD103+ IELs) were sorted into tubes containing 100μL TCL lysis buffer (Qiagen) supplemented with 1% β-mercaptoethanol, and stored at −80°C until cDNA synthesis (see **Figure S12A**). Total RNA was extracted using RNAclean XP beads (Agencourt) following the manufacture’s protocol. A modified SMART protocol (Quigley et al., 2011) was used in first-strand cDNA synthesis, described in detail elsewhere (Risnes et al., 2018). In brief, TCRα and TCRβ genes were amplified in two rounds of semi-nested PCR reactions. In the first reaction, the cDNA from each sample was divided in 3 replicates and amplified with *TRAC* and *TRBC* primers, and in the second PCR reaction different indexed primers were used to barcode the samples and replicates. A final third PCR reaction was performed using Illumina Seq Primers R1/R2 for sequencing. TCRα and TCRβ PCR products were quantified using the KAPA library quantification kit for Illumina platforms (KAPA Biosystems) and pooled at the same concentrations. Subsequently, pooled TCRα and TCRβ products were cleaned and concentrated with MinElute PCR Purification Kit (Qiagen), and ran on a 1.5% agarose gel. Bands of appropriate size (∼650bp) were gel-extracted (**Figure S12B**), purified using the Monarch Gel Extraction kit (New England Biolabs), and cleaned with MinElute PCR Purification Kit (Qiagen). The amplicon TCR library was sequenced using the paired-end 300 bp Illumina MiSeq and the approximate number of paired-end reads generated per CD8 T cell population was: 35,000 reads for TCRα and 45,000 reads for TCRβ sequences. The total number of reads was 60 million (approximately 20 million reads per library). Bulk TCR-seq data was preprocessed (VDJ alignment, clonotyping) using the MiXCR software package (Bolotin et al., 2015). Clonotypes were defined based on the V-gene, J-gene usage and the nucleotide sequence of the CDR3 region (Greiff et al., 2015b). Correlation of the read counts for the clonotypes found within the three molecular replicates was high **(Figure S12C)**, indicating reproducibility of the TCR repertoire sequencing (Greiff et al., 2014). For downstream analysis, raw reads from molecular triplicates were cumulated, and only clonotypes with a minimal read count of 10 were used. Sample preparation and read statistics are summarized in **Figure S12.** tcR package was used for descriptive statistics (Nazarov et al., 2015). The percentage of overlapping clones shared between two CD8 T cell subsets was calculated as: 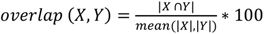, where |X| and |Y| are the clonal sizes (number of unique clones, either nucleotide or amino acid, see main text)) of repertoires X and Y.

### Statistical analysis

Statistical analyses were performed in Prism 7 (GraphPad Software). To assess statistical significance among SI CD8 T cell subsets, data were analyzed by one-way ANOVA (standard or repeated measures, RM-ANOVA) followed by Tukey’s multiple comparison tests. Replacement data and distribution of CD8 T cell subsets at different time points were analyzed by two-way ANOVA matching across subsets followed by Tukey’s multiple comparison tests. Correlations between replacement kinetics of different CD8 T cell subsets were calculated using Pearson correlation with two-tailed p-value (95% confidence interval). P-values of <0.05 were considered significant. TCR repertoire analysis was performed using the R statistical programming environment (R Development Core Team, 2018). Non-base R packages used for analyses were: tcR (Nazarov et al., 2015), upsetR (Lex et al., 2014), ggplot2 (Wickham, 2009), VennDiagram (Chen, 2018). Morisita-Horn index was calculated using the R divo package (Sadee et al., 2017). Vegan package (Oksanen et al., 2018) was used to calculate the diversity (Shannon-Evenness index) as previously described (Greiff et al., 2015a).

## Acknowledgments

We are grateful to Kjersti Thorvaldsen Hagen, Frank Sætre and Kathrine Hagelsteen for excellent technical assistance; the staff at the Endoscopy Unit and the surgery theatre; Christian Naper for providing HLA-typing; the Confocal Microscopy and Flow Cytometry Core Facilities at Oslo University Hospital – Rikshopitalet and Thomas S. Kupper (Dept. of Dermatology Harvard Medical School, Boston) for critical reading of the manuscript.

## Author contributions

R.B.C, O.J.B.L, E.S.B, and F.L.J. conceived the project. R.B.C., O.J.B.L, and S.K. processed samples, designed and performed experiments and analyzed data. L.R. helped establish methodology and advised on advanced flow-cytometry analysis. D.P, and L.F.R assisted in experiments. V.G. provided critical insights, assisted in bioinformatic analysis and design of figures, R.S.N., Y.Y. and V.G. developed bioinformatic tools. S.Y. and R. H. coordinated recruitment of patients and collection of biopsies. S.Y., R. H., O.M. Ø., and E.M.A. performed surgery and provided biopsies. V. P. performed endoscopy and provided endoscopic biopsies. R.B.C. performed bioinformatic analysis and prepared figures. R.B.C. and F.L.J. wrote the manuscript. O.J.B.L, S.K., V.G., L.R., D.P., L.F.R., E.M.A., S.W.Q, L.S., and E.S.B. contributed to writing the manuscript. S.W.Q, E.S. B, and F.L.J. supervised the study.

The authors declare no competing financial interests.

## Data and materials availability

Single cell and bulk TCR data: that data will be made available in SRA/ENA upon publication.

