## Supplemental materials_Bartolome et al. for "Resident memory CD8 T cells persist for years in human small intestine" (from another version of this preprint: version 1, 2019-02-18)

**Supplementary Materials**

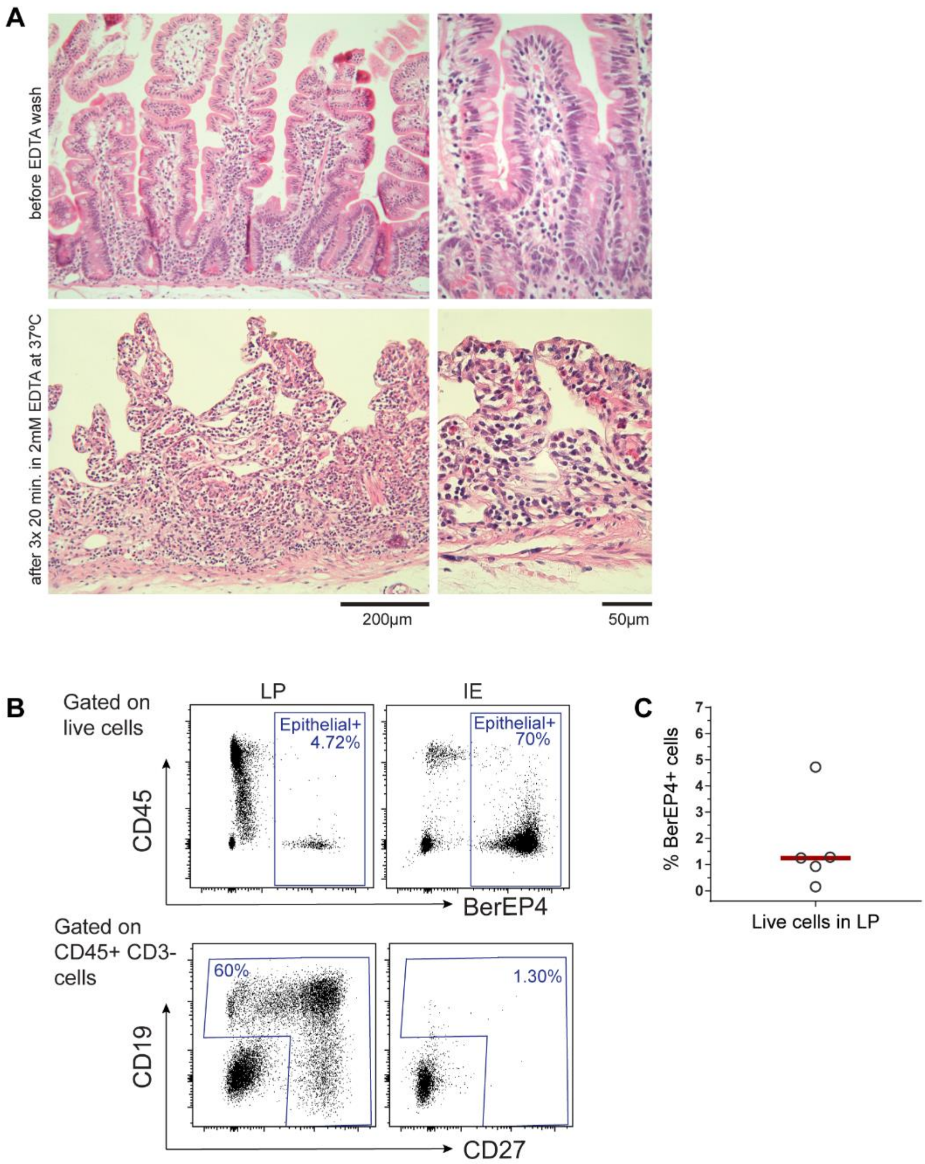


**Figure S1.** Confirmation of the absence of cross-contamination between LP and IE tissue fractions. (**A**) Representative hematoxylin-eosin (H/E) staining of tissue sections obtained before and after three sequential washing steps with EDTA buffer (PBS containing 2mM EDTA and 1%FCS) at 37°C with vigorous shaking. (**B**) Representative flow cytometric plot showing the percentage of epithelial cells (BerEP4+ cells) and B cells within each fraction. (**C**) Compile data for the percentage of epithelial cells (BerEP4+ cells) remaining in the LP after three sequential washing steps with EDTA buffer.

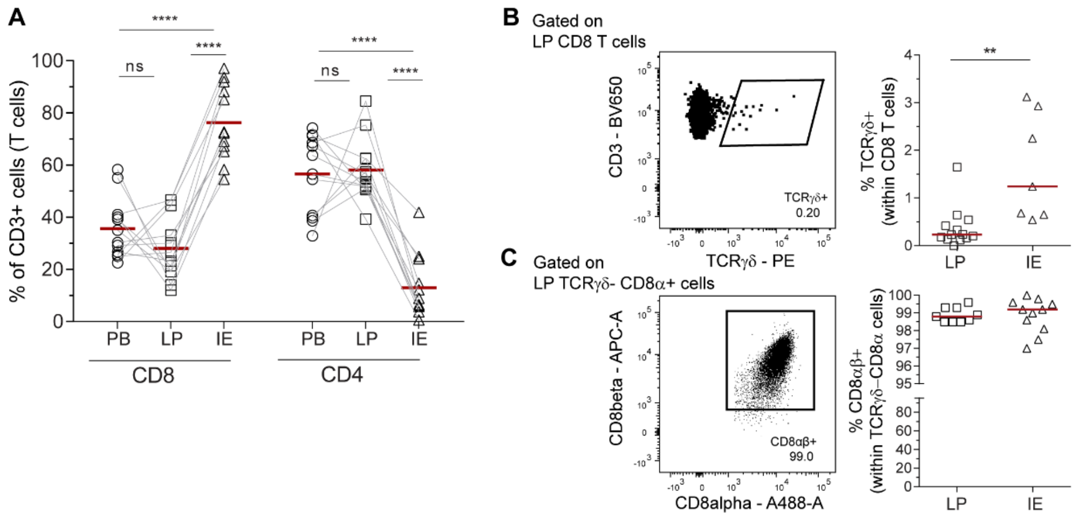


**Figure S2**. (**A**). Percentage of T cell subsets (CD4 and CD8) in peripheral blood (PB), lamina propria (LP) and epithelium(IE) measured by flow cytometry. Red line indicates mean value. Statistics performed using two-ways ANOVA, repeated measures (RM) matching both factors, and Turkey's multiple comparison test. (**B**) TCRγδ and (**C**) CD8β expression on LP and IE CD8 T cells analyzed by flow cytometry. Red line indicates median value. Unpaired T-test. **P≤0.01.

11 colors - 3 compartments (PB+ LP+ IE) FACS analysis

| | 405 nm | | | 488 nm | | 561 nm | | | 640 nm | | |
|---|---|---|---|---|---|---|---|---|---|---|---|
| | eF450 | BV605 | BV650 | FITC | PerCP-Cy5.5 | PE | PE-CF594 | PE-Cy7 | APC | Ax700 | APC H7 |
| Tube1 | CD4 | PD1 | CD3 | CD103 | CD8 | KLRG1 | CCR7 | CD45-RA | CD69 | CD45 | L/D |
| Tube2 | CD4 | CD127 | CD3 | CD103 | CD8 | KLRG1 | CCR7 | CD45-RA | CD28 | CD45 | L/D |
| Tube3 | CD4 | CD161 | CD3 | CD103 | CD8 | KLRG1 | CCR7 | CD45-RA | CD244 | CD45 | L/D |
| Tube4 | CD4 | NKG2D | CD3 | CD103 | CD8 | KLRG1 | CCR7 | CD45-RA | CD45-RO | CD45 | L/D |

x3 = 12 tubes
PB
LP
IE

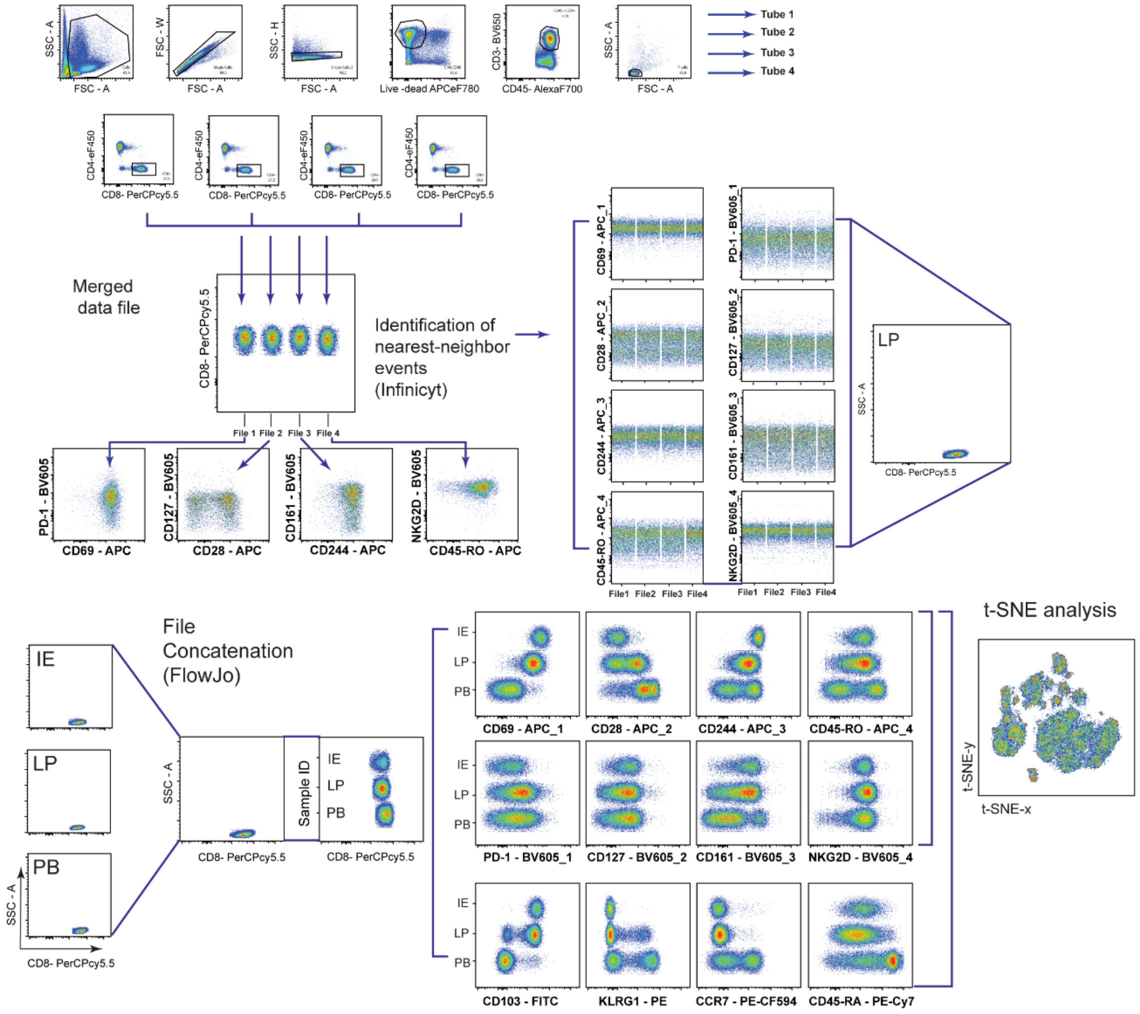


**Figure S3**. Related to Figure 1C. Panel design, merge (as described in (Pedreira et al., 2008a; Pedreira et al., 2008b) ) and concatenation of flow cytometric files before applying t-SNE analysis (FlowJo plugin).

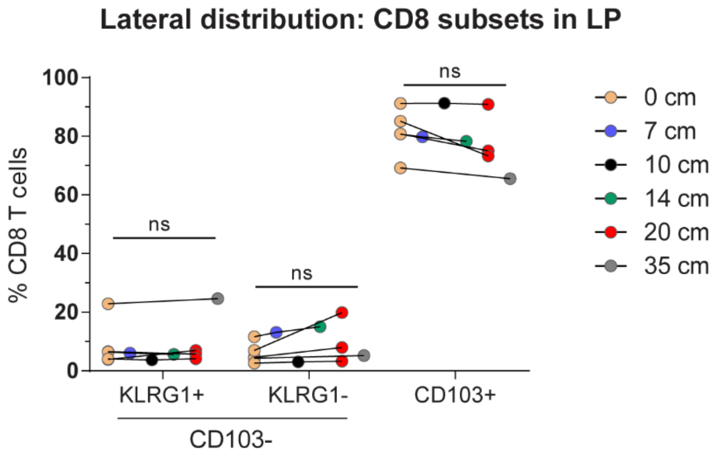


**Figure S4.** Lateral distribution of LP CD8 T cell subsets. The lengthwise representation of the CD8 subsets in LP was determined by flow cytometric analysis of biopsies taken at intervals along resected duodenum-proximal jejunum from individual subjects after Whipple procedure. n = 5; paired Student's t test comparing 0 cm –furthest distance. ns, not significant.

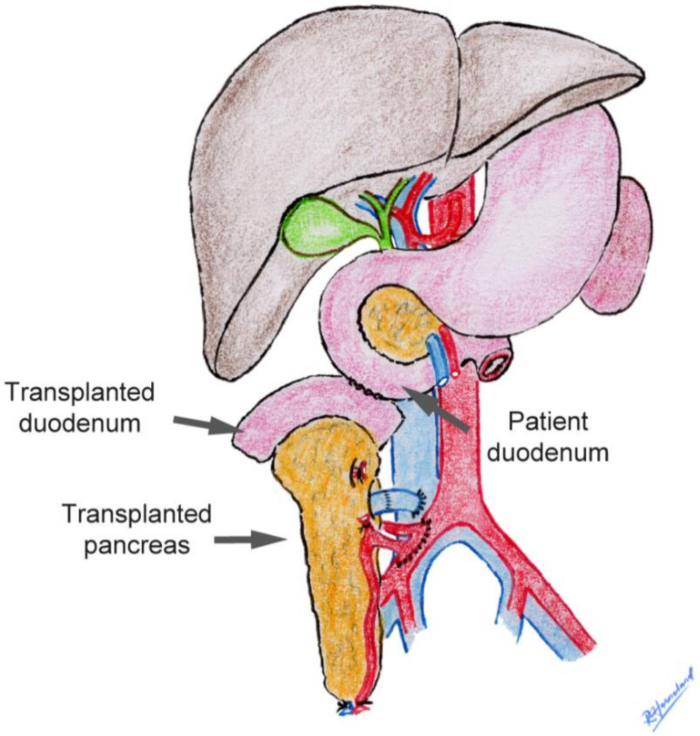


**Figure S5.** Representation of the pancreas transplantation showing the duodeno-duodenostomy anastomosis, as described in (Horneland et al., 2015).

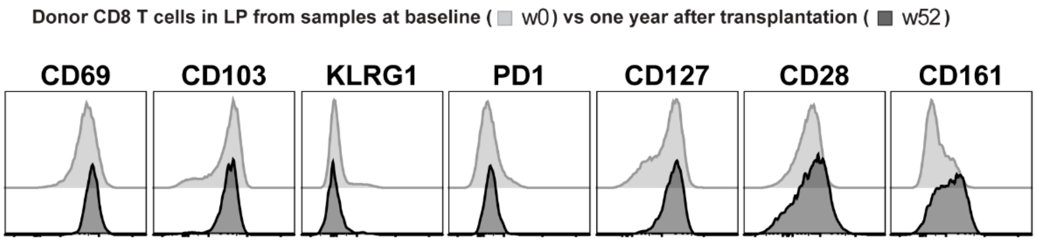


**Figure S6.** The expression profile of donor LP CD8 T cells derived from duodenal transplant at baseline (w0) vs one-year after transplantation (w52).

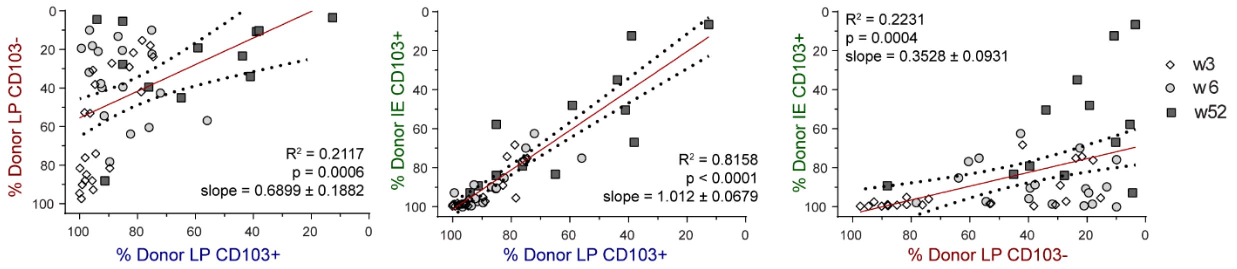


**Figure S7.** Correlation of replacement kinetics for donor CD103-, LP and IE CD103+ CD8 T cell subsets. Statistics performed using Pearson correlation with two-tailed p-value (95% confidence interval).

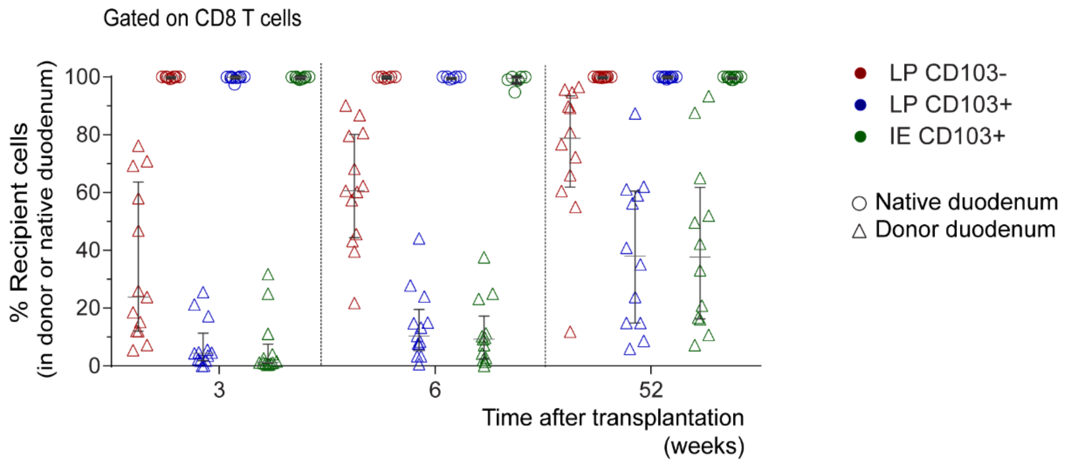


**Figure S8.** Percentage of recipient CD8 T cells for each subset isolated from donor and recipient (native) duodenum of the same patients at different time points post-transplantation.

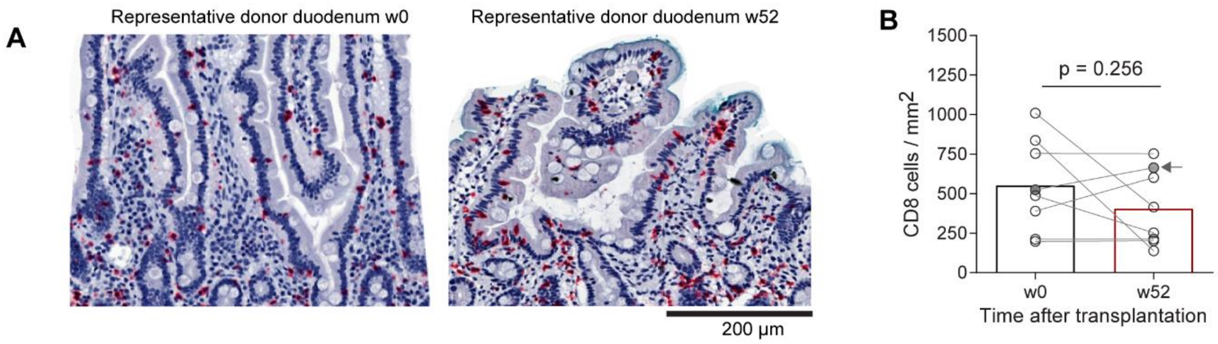


**Figure S9. (A)** Representative immunohistochemistry staining of CD8 T cells (Fast-Red) on tissue sections from donor duodenum of a representative patient before (w0) and 1-year post-Tx (w52). **(B)** Total CD8 T cell counts on tissue sections from donor duodenum of representative patients (n=8) before (w0) and 1-year post-Tx (w52). The patient shown in (A) is filled in gray and pointed with an arrow. Paired T-test.

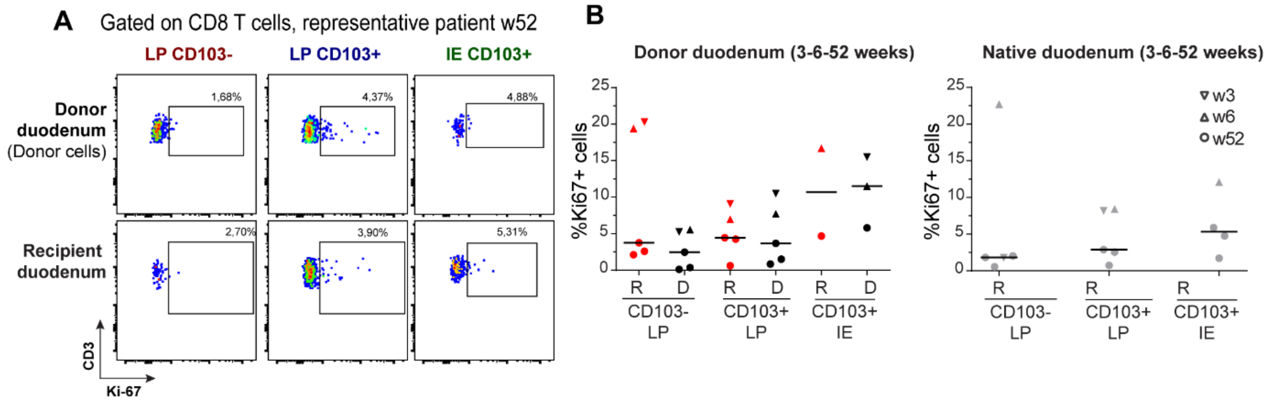


**Figure S10. (A)** Representative dot plot of Ki67 expression in CD8 T cell subsets derived from a donor and native duodenum one-year post-transplantation. **(B)** Compiled data for the percentage of CD8 T cell subsets expressing Ki67 at different time points after transplantation in donor and native duodenum.

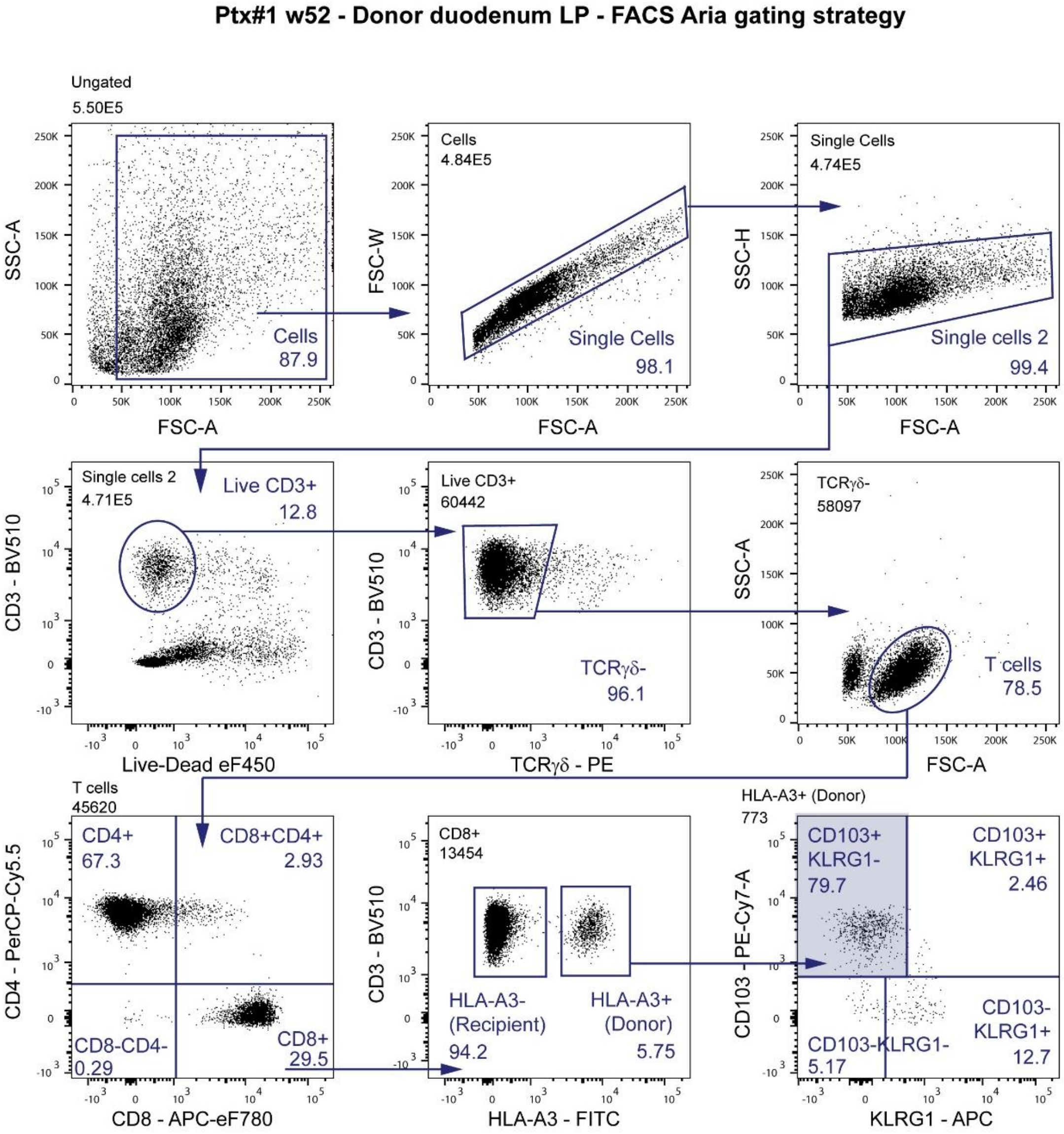


**Figure S11**. Gating strategy for single cell sorting of CD8 T cells from duodenal biopsies. Representative sample of grafted duodenum from a transplanted patient one-year after transplantation is shown.

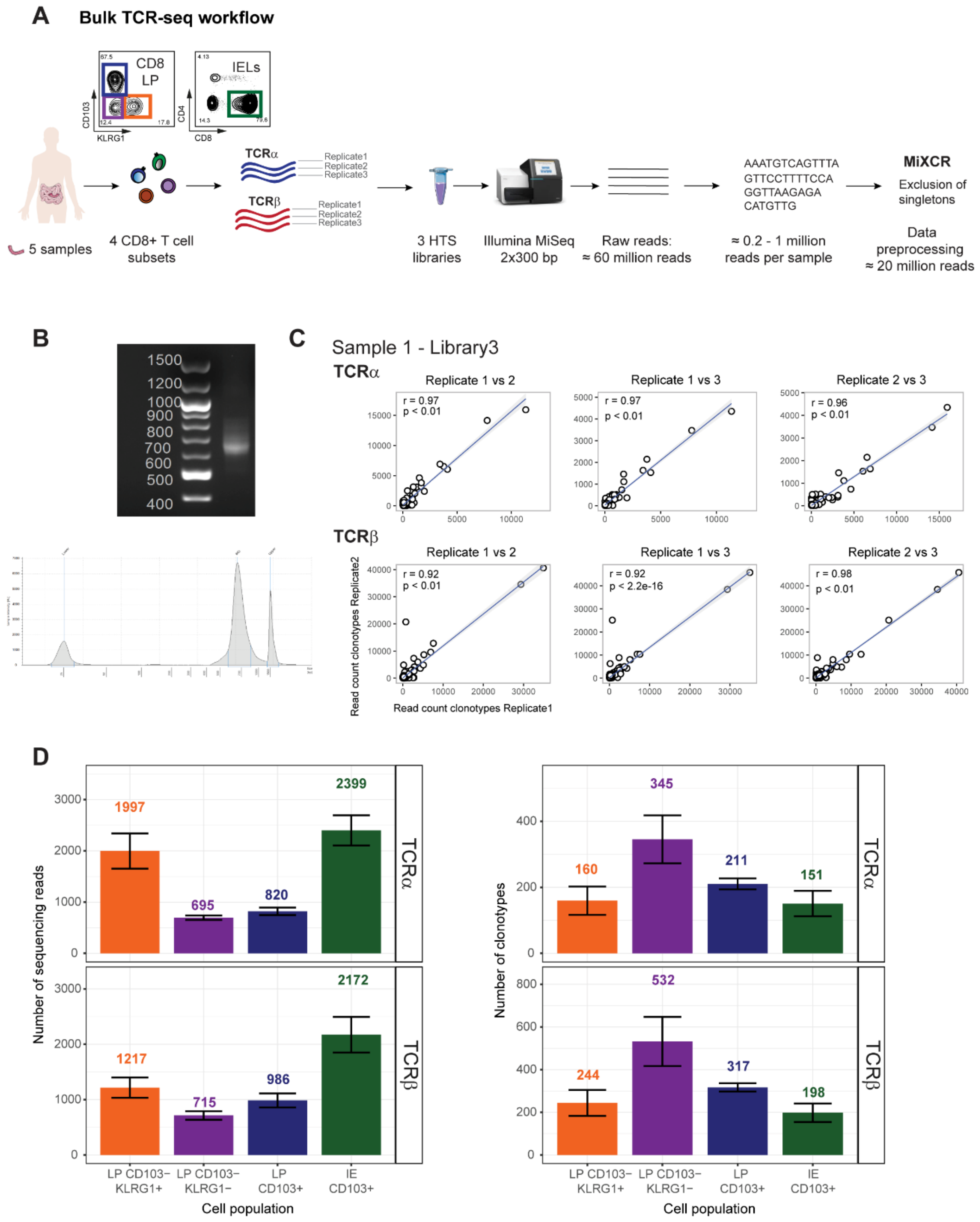


**Figure S12.** Bulk TCR-seq workflow and read statistics. (**A**) Overview of sample processing, data acquisition and data processing. (**B**) Agarose gel shows representative library following the protocol explained in (A). Expected band size ≈700 bp. (**C**) Correlation plots showing the correlation of the common clonotypes between the three replicates representative sample. (**D**) Clone and sequence counts after MiXCR preprocessing are shown by CD8 T cell subset (mean ± s.e.m). The total number of sequencing reads used for the analysis was 20,692,061 (total number of raw reads: ≈ 62,317,193). Mean Phred scores of raw data were ≥30.

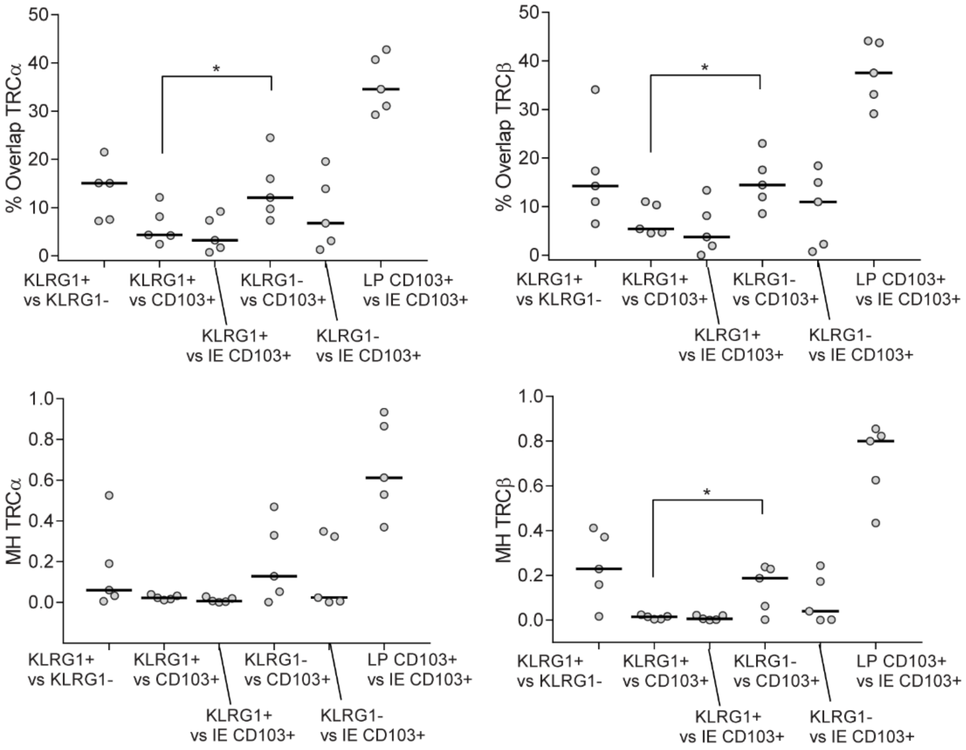


**Figure S13.** Compiled data of pairwise clonal overlap and Morisita-Horn calculation for the TCRα and TCRβ clonotypes in all the samples. Paired t-test, two-tailed; *P≤0.05, n.s., not significant.

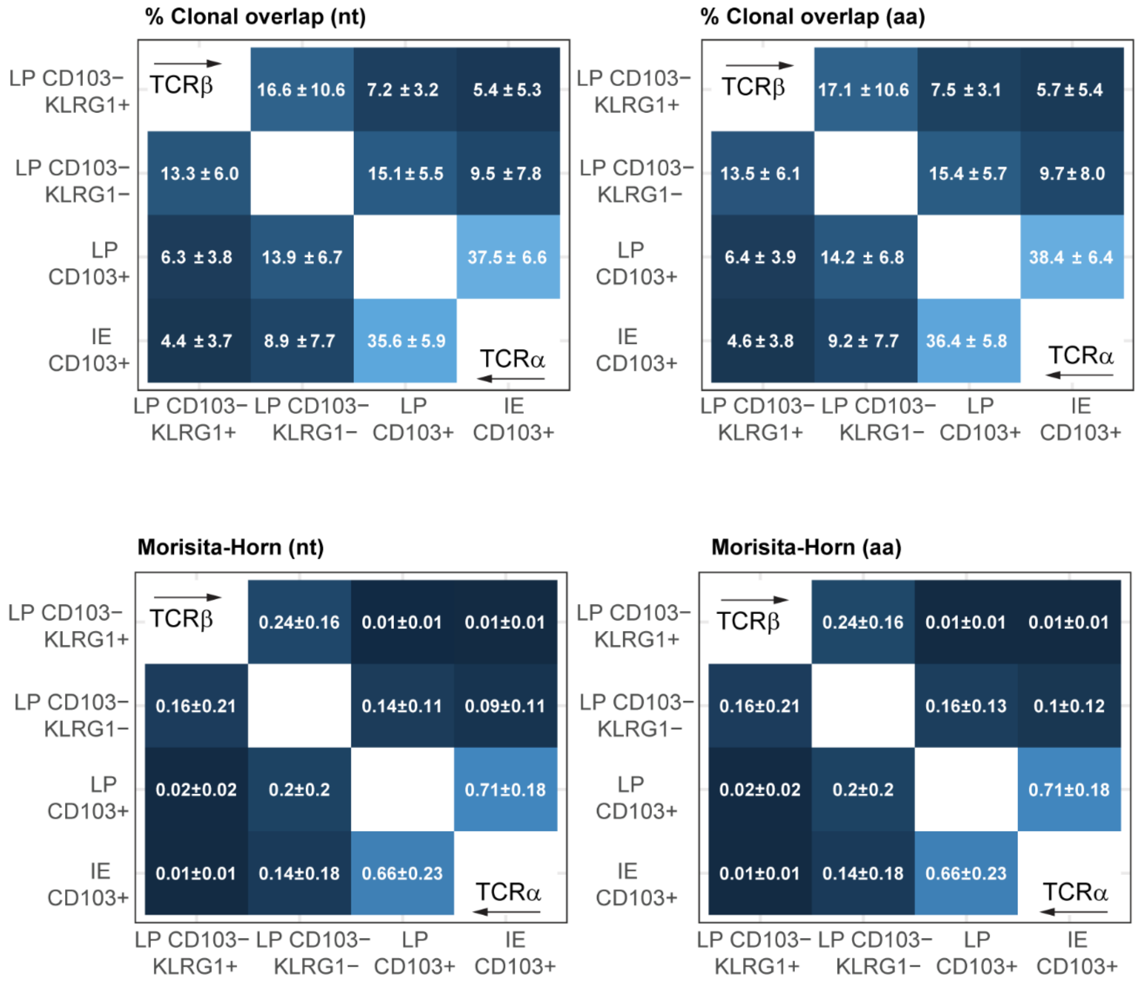


**Figure S14.** Pairwise clonal overlap and Morisita-Horn calculation for the TCRα and TCRβ clonotypes in all the samples. Similar values were found when the amino acid sequence (aa) of the CDR3 was used as the clonotype definition instead of the nucleotide sequence (nt).

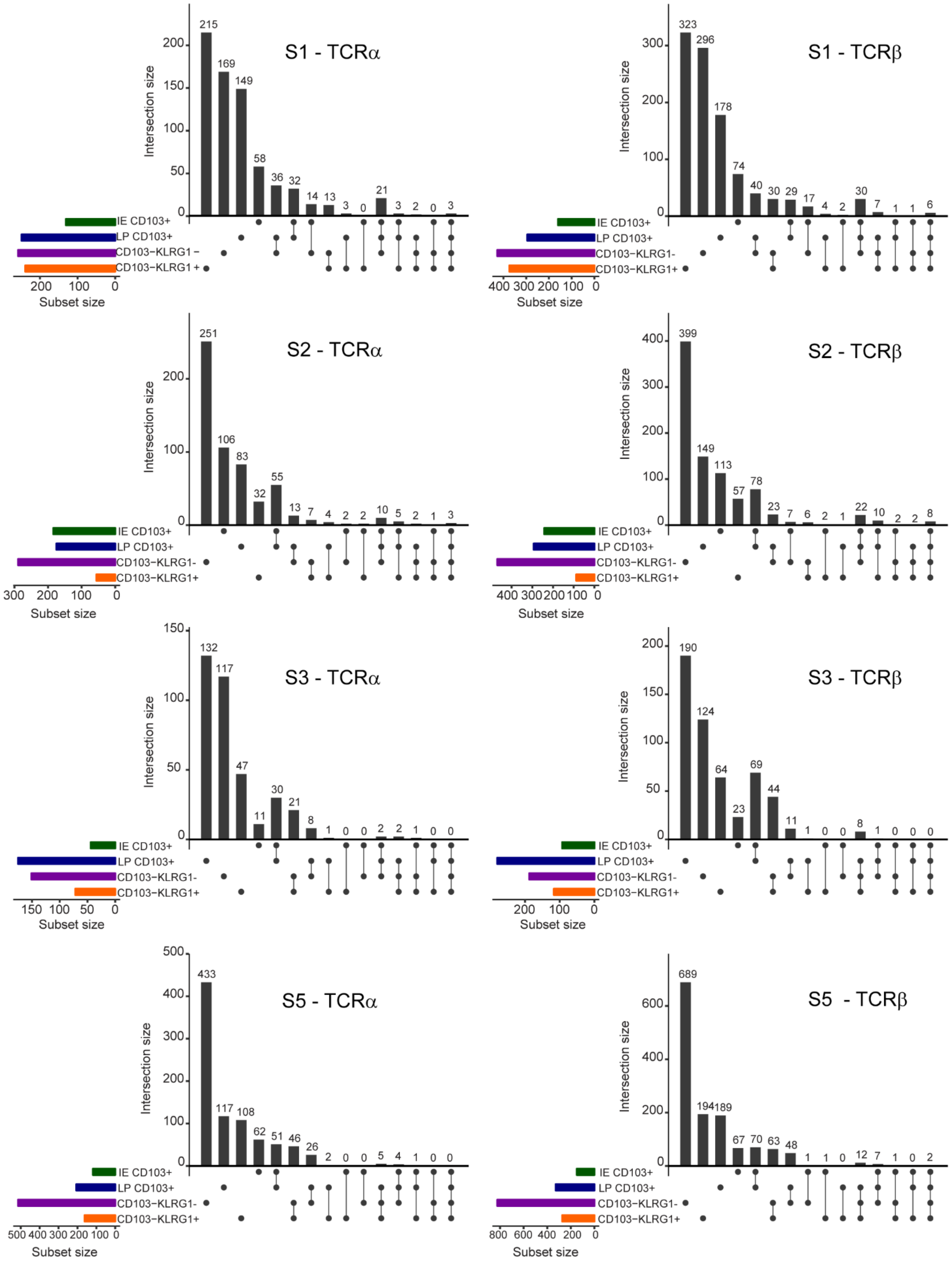


**Figure S15.** Upset plots with the overlapping clones for TCRα and TCRβ in all the samples analyzed.

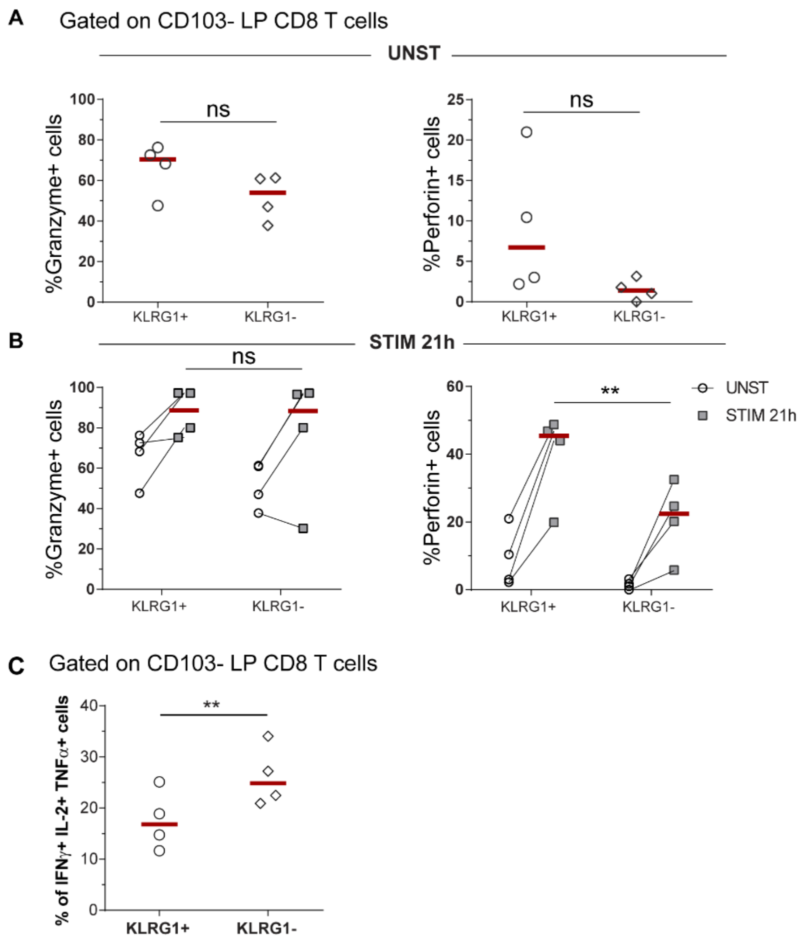


**Figure S16.** Percentage of Granzyme and perforin positive cells in unstimulated samples (**A**) and after 21h of activation with anti-CD3 beads (**B)** for the different CD103- CD8 T cell subsets (KLRG1+, KLRG1-) in LP. (**C)** Percentage of polyfunctional (IL-2+, IFN-γ+ and TNF-α+) cells for the different CD103- CD8 T cell subsets (KLRG1+, KLRG1-) in LP.

**Table S1.** List of antibodies used in the study

| Target | Clone | Fluorophore | Company | Cat. num | Phenotype Trm | PTx panel | Aria Sorting | Ki67 nuclear | Cytotoxicity | Cytokines | X/Y - FISH |
|---|---|---|---|---|---|---|---|---|---|---|---|
| CD3 | OKT_3 | BV650 | Biolegend | 317324 | x | x | | x | | | |
| CD3 | OKT_3 | APC-eF780 | eBioscience | 47-0037-42 | | | x | | | x | |
| CD3 | OKT_3 | BV510 | Biolegend | 317332 | | | | | | x | |
| CD3 | OKT_3 | APC | Biolegend | 317318 | | | x | | | | |
| CD3 | OKT_3 | PerCP-Cy5.5 | Biolegend | 317336 | | | | x | | | |
| CD3 | Poly | unconjugated | Dako | A0452 | | | | | | | x |
| CD4 | OKT_4 | eF450 | eBioscience | 48-0048-42 | x | x | | x | | x | |
| CD4 | OKT_4 | PerCP-Cy5.5 | Biolegend | 317428 | | | x | | | | |
| CD8 | SK1 | Alexa-Fluor488 | Biolegend | 344716 | x | x | | | | | |
| CD8 | SK1 | APC-eF780 | eBioscience | 47-0087-42 | | | x | | x | x | |
| CD8 | SK1 | PerCP Cy5.5 | Biolegend | 344710 | x | | | | | x | |
| CD8 | SK1 | PE | BD Biosciences | 340046 | | | | x | | | |
| CD8 | 4B11 | unconjugated | Novocastra | NCL-L-CD8-4B11 | | | | | | | x |
| CD8b /NHP | SIDI8BEE | eF660 | eBioscience | 50-5273-41 | x | x | | | | | |
| CD28 | CD28.2 | BV605 | BD Horizon | 562976 | x | x | | | | | |
| CD28 | CD28.2 | APC | Biolegend | 302912 | x | x | | | | | |
| CD28 | CD28.2 | PE | Biolegend | 302908 | x | x | | | | | |
| CD45 | HI30 | BV510 | Biolegend | 304036 | | | x | x | x | | |
| CD45 | HI30 | Ax700 | Biolegend | 304024 | x | | | | | | |
| CD45 | 2D1 | APC-H7 | BD-Biosciences | 560178 | x | x | | | | x | |
| CD45-RA | HI100 | APC-eF780 | eBioscience | 47-0458-42 | x | | | | | x | |
| CD45-RA | HI100 | PE-Cy7 | eBioscience | 25-0458-42 | | | x | | | x | |
| CD45-RO | UCHL1 | APC | eBioscience | 17-0457-42 | | | | | | | |
| CD103 | B-Ly7 | PE-Cy7 | Biolegend | 350212 | x | x | x | x | x | | |
| CD103 | Ber-ACT8 | BV605 | Biolegend | 350218 | | | | | | x | |
| CD103 | Ber-ACT8 | PE-Cy7 | Biolegend | 350212 | | | | | | x | |
| CD103 | B-Ly7 | FITC | eBioscience | 11-1038-42 | x | | | | | | |
| CD62-L (L-Sel) | SK11 | FITC | BD Pharmingen | 347443 | x | | | | | | |
| CD127 (IL7-R) | Hil7r-m21 | PE | BD Pharmingen | 561028 | x | x | | | | x | |
| CD127 (IL7-R) | Hil7r-m21 | BV605 | BD Horizon | 562662 | x | x | | | | | |
| CD161 (KLRB1) | HP-3G10 | BV605 | Biolegend | 339915 | x | x | | | | | |
| CD244 (SLAMF4) | 2B4 | APC | Biolegend | 329511 | x | x | | | | | |
| CCR7 (CD197) | G043H7 | PE | Biolegend | 353203 | x | | | | | | |
| CCR7 (CD197) | G043H7 | PE-dazzle594 | Biolegend | 353235 | x | | | | | | |
| PD-1 (CD279) | EH12.1 | BV605 | BD Horizon | 563245 | x | x | | | | | |
| KLRG1 | 13F12F2 | APC | eBioscience | 17-9488-42 | x | x | x | x | x | x | |
| KLRG1 | 13F12F2 | PE | eBioscience | 12-9488-42 | x | x | x | | x | | |
| NKG2D | 1D11 | PE | Biolegend | 320805 | x | x | | | | | |
| NKG2D | 1D11 | BV605 | BD Biosciences | 743559 | x | x | | | | | |
| TCR-gd | 5A6.E9 | PE | Molecular Probes™, Invitrogen | MHGD04 | x | x | x | | | | |
| TCR-gd | 5A6.E9 | FITC | Molecular Probes™, Invitrogen | MHGD01 | | x | x | | | | |
| Anti-Human Epithelial Antigen | Ber-EP4 | FITC | Dako | F0860 | | | x | | | | |
| HLA-A2 | BB 7.2 | PE | Abcam | ab79523 | | x | | | | | |
| HLA-A3 | GAP.A3 | FITC | eBioscience | 11-5754-42 | | x | | | | | |
| HLA-A3 | GAP.A3 | APC | eBioscience | 17-5754-42 | | x | x | | | | |
| HLA-B7 | BB7.1 | PE | Millipore | MAB1288 | | x | | | | | |
| HLA-B8 | REA145 | PE | Miltenyi Biotech | 130-118-960 | | x | x | | | | |
| Granzyme-B | CLB-GB11 | PE | SANQUIN (Dianova AS) | M2289 | | | | | x | | |
| Perforin | gG9 | FITC | BD Pharmingen | 556577 | | | | | x | | |
| TNF-alpha | MAb11 | APC | Biolegend | 502912 | | | | | | x | |
| IFNg | 4S.B3 | Ax488 | Biolegend | 502515 | | | | | | x | |
| IL2 | MQ1-17H12 | PE | Biolegend | 500307 | | | | | | x | |
| IL2 | MQ1-17H12 | BV421 | Biolegend | 500327 | | | | | | x | |
| MIP1beta (CCL4) | FL3423L | APC | eBioscience | 17-7540-41 | | | | | | x | |
| Ki67 | B56 | Ax488 | BD Pharmingen | 558616 | | | | x | | | |
| Isotype mouse IgG1 | MOPC-21 | Ax488 | BD Pharmingen | 555909 | | | | x | x | | |
| Isotype mouse IgG2b | 27-35 | FITC | BD Pharmingen | 556577 | | | | | x | | |
| Isotype mouse IgG1 | MOPC-31C | PE | BD Pharmingen | 550617 | | | | | x | | |
| Mouse IgG2b | Poly | Alexa 555 | Molecular Probes™, Invitrogen | A-21147 | | | | | | | x |
| Rabbit IgG (H+L) | Poly | Alexa 647 | Molecular Probes™, Invitrogen | A-31573 | | | | | | | x |
